# A common mechanism of Sec61 translocon inhibition by small molecules

**DOI:** 10.1101/2022.08.11.503542

**Authors:** Samuel Itskanov, Laurie Wang, Tina Junne, Rumi Sherriff, Li Xiao, Nicolas Blanchard, Wei Q. Shi, Craig Forsyth, Dominic Hoepfner, Martin Spiess, Eunyong Park

**Affiliations:** Biophysics Graduate Program, University of California, Berkeley, Berkeley, CA 94720, USA; Department of Molecular and Cell Biology, University of California, Berkeley, CA 94720, USA; Biozentrum, University of Basel, CH-4056, Basel, Switzerland; Department of Chemistry and Biochemistry, The Ohio State University, Columbus, Ohio 43210, United States; CNRS, LIMA, UMR 7042, Université de Haute-Alsace, Université de Strasbourg, Mulhouse, France; Department of Chemistry, Ball State University, Muncie, IN 47306, USA; Novartis Institutes for BioMedical Research, Novartis Pharma AG, Forum 1 Novartis Campus, CH-4056, Basel, Switzerland; California Institute for Quantitative Biosciences, University of California, Berkeley, CA 94720, USA

## Abstract

The Sec61 complex forms a protein-conducting channel in the endoplasmic reticulum (ER) membrane that is required for secretion of soluble proteins and production of many membrane proteins. Several natural and synthetic small molecules specifically inhibit the Sec61 channel, generating cellular effects that are potentially useful for therapeutic purposes, but their inhibitory mechanisms remain unclear. Here we present near-atomic-resolution structures of the human Sec61 channel inhibited by a comprehensive panel of structurally distinct small molecules— cotransin, decatransin, apratoxin F, ipomoeassin F, mycolactone, cyclotriazadisulfonamide (CADA) and eeyarestatin I (ESI). Remarkably, all inhibitors bind to a common lipid-exposed pocket formed by the partially open lateral gate and plug domain of the channel. Mutations conferring resistance to the inhibitors are clustered at this binding pocket. The structures indicate that Sec61 inhibitors stabilize the plug domain of Sec61 in a closed state, thereby preventing the protein-translocation pore from opening. Our study reveals molecular interactions between Sec61 and its inhibitors in atomic detail and offers the structural framework for further pharmacological studies and drug design.

## Introduction

The universally conserved heterotrimeric Sec61 complex (SecY in prokaryotes) plays essential roles in biosynthesis of more than one third of proteins in all species (for review, see ref. ^1-4^). In eukaryotes, secretory proteins are first translocated into the ER by the Sec61 complex before reaching the cell surface by vesicular trafficking. The Sec61 complex also mediates membrane integration of many proteins, including most cell surface receptors and cell adhesion molecules. The Sec61/SecY channel has an hourglass-like structure with a pore constriction (termed the pore ring) halfway across the membrane, which is gated by a movement of a plug-like ER-lumenal (or extracellular in SecY) domain of the channel^5^. In addition, the channel has a seam (lateral gate) in the wall that can open laterally in the plane of the membrane to release transmembrane segments (TMs) of membrane protein clients into the lipid phase. Concerted opening of the lumenal and lateral gates is also required for initial insertion of the client protein’s hydrophobic signal sequence or anchor into the channel (Fig. 1a).

**Figure 1.**
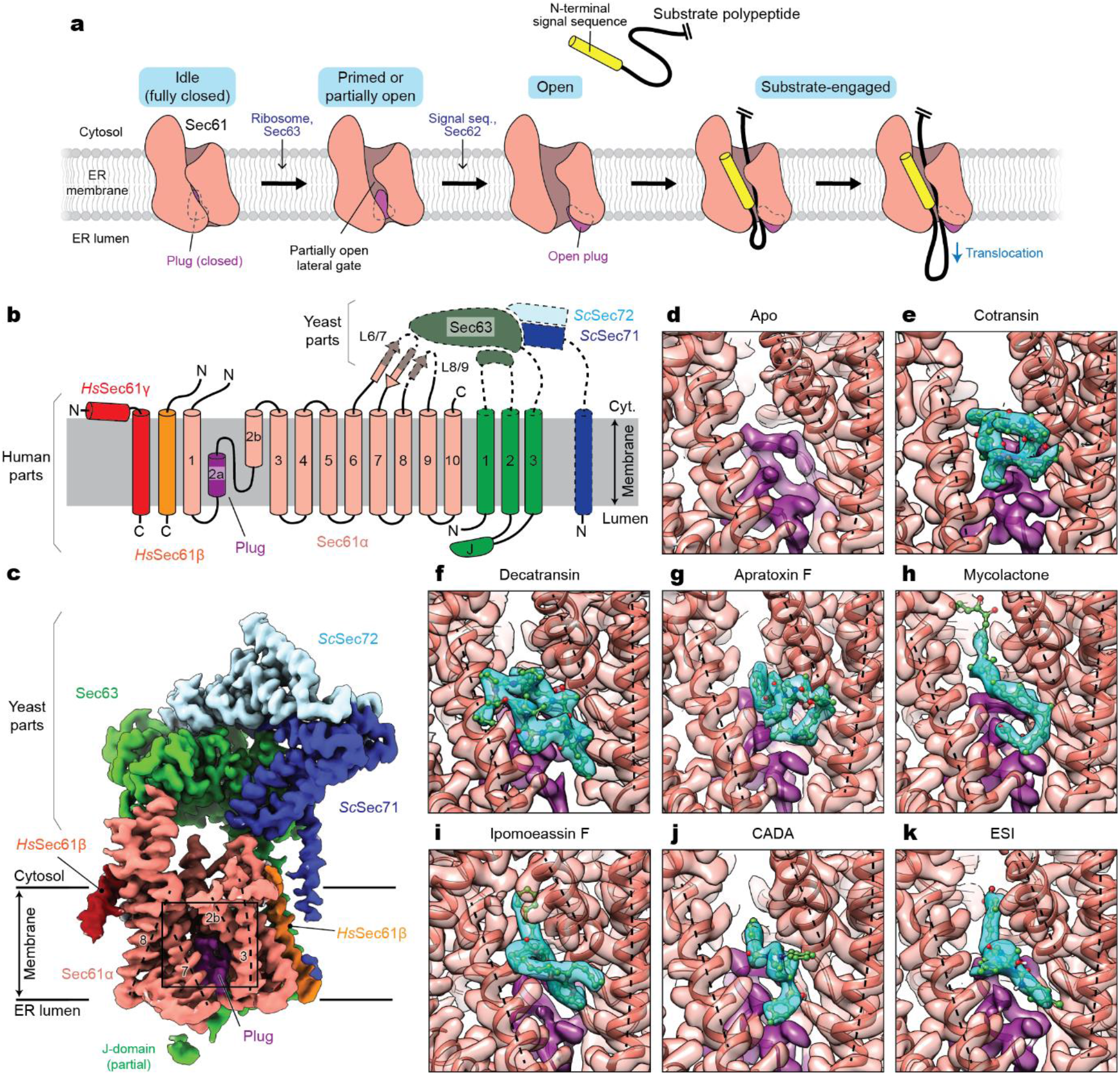
Cryo-EM structures of the human Sec61 complex inhibited by various small-molecule inhibitors. **a**, Architecture of the Sec61 channel and overall model for gating and substrate engagement. **b**, Design of a human-yeast chimeric Sec complex. Parts derived from human and yeast proteins are outlined with solid and dashed lines, respectively. Note that except for the cytosolic L6/7 and L8/9 loops, Sec61α is from the human sequence (SEC61A1). *Hs, Homo sapiens*; *Sc, Saccharomyces cerevisiae*; J, J-domain. **c**, 2.7-Å-resolution cryo-EM map of the chimeric Sec complex in an apo state (Class 1, unsharpened map). The lateral gate helices are indicated by dashed lines and TM numbers. The region outlined by a rectangle indicates the inhibitor-binding site (also see **d**–**k**). **d**–**k**, Views into the inhibitor-binding site of Sec61α of apo and inhibitor-bound structures. Cryo-EM maps (semi-transparent surface) and atomic models were overlaid. Inhibitor and plug densities are shown in cyan and purple, respectively. Dashed lines indicate lateral gate helices TMs 2b, 3, and 7 as in c.

The Sec61/SecY channel translocates polypeptides either co-translationally by docking a translating ribosome or post-translationally by engaging a fully synthesized polypeptide client. In eukaryotes, the post-translational mode is enabled by association of the channel with two additional membrane proteins Sec63 and Sec62 (ref. ^6-8^). X-ray crystallography and cryo-electron microscopy (cryo-EM) have visualized structures of the Sec61/SecY channel in different functional states and revealed how it is gated and engages with client proteins^5,9-18^. The current model posits that association of a ribosome or Sec63 slightly perturbs (“primes”) or partially opens the lateral gate^11,15,16^ (Fig. 1a). Insertion of the client polypeptide needs further widening of the lateral opening and a displacement of the plug away from the pore, which occur in a cooperative manner. In cotranslational translocation, these conformational changes are presumed to be induced by an interaction between the channel and the signal sequence/anchor^11,13^, whereas in post-translational translocation, they seem to be mediated by Sec62^17^.

Several natural and synthetic small molecules bind to Sec61 and inhibit protein translocation (for review, see ref. ^19-22^). These inhibitors have been investigated as potential anticancer, antiviral, and/or immunosuppressive agents^23-27^. Inhibition of Sec61 leads to downregulation of disease-related and clinically-relevant proteins, such as cytokines, cell surface receptors, and viral membrane proteins. Indeed, one such Sec61 inhibitor is currently being tested in a phase-I clinical trial for treatment of solid tumor malignancies^28^. A founding class of Sec61 inhibitors is a group of fungal-derived cyclic heptadepsipeptides named cotransins^29-31^. Other naturally occurring inhibitors discovered to date are decatransin, mycolactone, apratoxins, coibamide A, and ipomoeassin F, which are produced by certain fungal, bacterial, and plant species^32-38^. In addition, two synthetic compounds CADA and ESI have also been shown to inhibit the Sec61 channel^39,40^. These inhibitors are structurally unrelated to each other, but several of them have been suggested to bind to an overlapping site in the Sec61 channel based on their abilities to compete for Sec61 binding. Remarkably, cotransin and CADA inhibit Sec61 in a client-specific manner^29,30,41^, whereas other inhibitors act more broadly independent of clients. Biochemical data suggest that cotransin likely interacts with the lateral gate and/or the plug of Sec61 (ref. ^42^). However, key information regarding the actions of these inhibitors remains unavailable, including molecular details about Sec61-inhibitor interactions, which specific steps along the translocation process are inhibited, and what underlies client-specific versus broad-spectrum inhibition. This has limited our capability to design or discover additional therapeutically promising small-molecule agents that target Sec61.

## Results and Discussion

### Experimental design and cryo-EM analysis of inhibitor-bound Sec61

To understand the mechanism of Sec61 inhibition, we sought to determine high-resolution structures of inhibitor-bound Sec61 using cryo-EM. To date, all mammalian Sec61 structures have been obtained from ribosome-bound cotranslational complexes^11,12,43^. However, due to flexibility of Sec61 with respect to the ribosome, this approach limits the resolution of Sec61 to only ∼5 Å, a resolution that is impractical to model protein side chains and small ligands^11^. This problem also exists in a recent cryo-EM structure of a mycolactone-treated Sec61-ribosome complex^44^. By contrast, we previously attained 3.1–3.7-Å resolution structures of the Sec61 channel from fungal post-translational translocation complexes^15,17^ (termed the Sec complex), which contained Sec62, Sec63 and fungal-specific nonessential Sec71 and Sec72 in addition to the three (α, β, and γ) subunits of Sec61. Thus, we reasoned that use of the Sec complex would be an effective approach to study Sec61 inhibitors.

To enable high-resolution cryo-EM analysis of inhibitor-bound human Sec61, we designed a chimeric Sec complex, whose transmembrane and cytosolic domains are derived from the human and yeast proteins, respectively (Fig. 1b). Our initial efforts employing the entirely yeast or human Sec complex were unsuccessful. The yeast Sec complex incubated with cotransin failed to show any cotransin-like feature in the cryo-EM map (Supplementary Fig. S1 a and b). This could be due to a lower binding affinity of cotransin towards yeast Sec61 compared to mammalian Sec61^32^, the presence of detergent in the sample, or both. While we could see a putative cotransin density in a cryo-EM structure of the human Sec complex lacking Sec62, the resolution could not be improved beyond ∼5 Å, probably due to high flexibility of the cytosolic domain of human Sec63 (Supplementary Fig. S1 c–f). On the other hand, the human-yeast chimeric Sec complex reconstituted into a peptidisc^45^ yielded structures at overall 2.5 to 2.9-Å-resolution with most side-chain densities well defined (Fig 1c, and Supplementary Figs. S2–S4 and Table S1). In the absence of inhibitors, particle images could be sorted into two three-dimensional (3-D) classes with minor differences (Supplementary Fig. S2 b–g). In both classes, the Sec61 channel adopts a similar conformation, including a partially open lateral gate and a closed plug, as expected for a complex lacking Sec62 (ref. ^17^). However, the two classes showed slightly different arrangements of Sec61 with respect to Sec63-Sec71-Sec72 due to a loose contact between the engineered L6/7 loop of Sec61α and the FN3 domain of yeast Sec63 in Class 2 (Supplementary Fig. S2 g and h).

For inhibitor-bound structures, we used five naturally occurring inhibitors, cotransin, decatransin, apratoxin F, ipomoeassin F, and mycolactone; and two designed synthetic compounds CADA and ESI. Focused refinement masking out the cytosolic domains of Sec63-Sec71-Sec72 further improved the map of the Sec61 complex (at overall resolution of 2.6 to 3.2 Å) showing clear, well-defined density features for the added inhibitor (Fig. 1 d–k, and Supplementary Figs. S3 and S4). Local resolution around the inhibitor-binding region was on par with or better than the overall resolution owing to relatively uniform resolution distributions (Supplementary Fig. S3f). Reliable atomic models of inhibitor molecules could be built into the densities of inhibitors based on their two-dimensional (2-D) chemical structures (Fig. 1 d–k). However, we note that positions and orientations of certain atoms and bonds may deviate from their true structures as our structures do not resolve individual atoms.

### Inhibitor-binding site

Despite their diverse chemical structures, all analyzed inhibitors are found to bind essentially to the same site in the Sec61 channel (Figs. 1 and 2, and Supplementary Fig. S5). The pocket is formed at the partially open lateral gate, approximately halfway across the membrane. The inhibitors commonly interact with lateral gate helices TMs 2b, 3, and 7 of the Sec61α subunit. However, it should be noted that the actual structure of the pocket substantially varies depending on the bound inhibitor because the lateral gate adopts different degrees of opening (Fig. 2, and Supplementary Fig. S5). The width of the lateral gate opening is widest in the cotransin-bound structure and narrowest in the ipomoeassin F-bound structure. During protein translocation, the lateral gate of the Sec61/SecY channel dynamically adopts closed or variable open states by a relative motion between the N- and C-terminal halves of the α subunit^5,9-18^.

**Figure 2.**
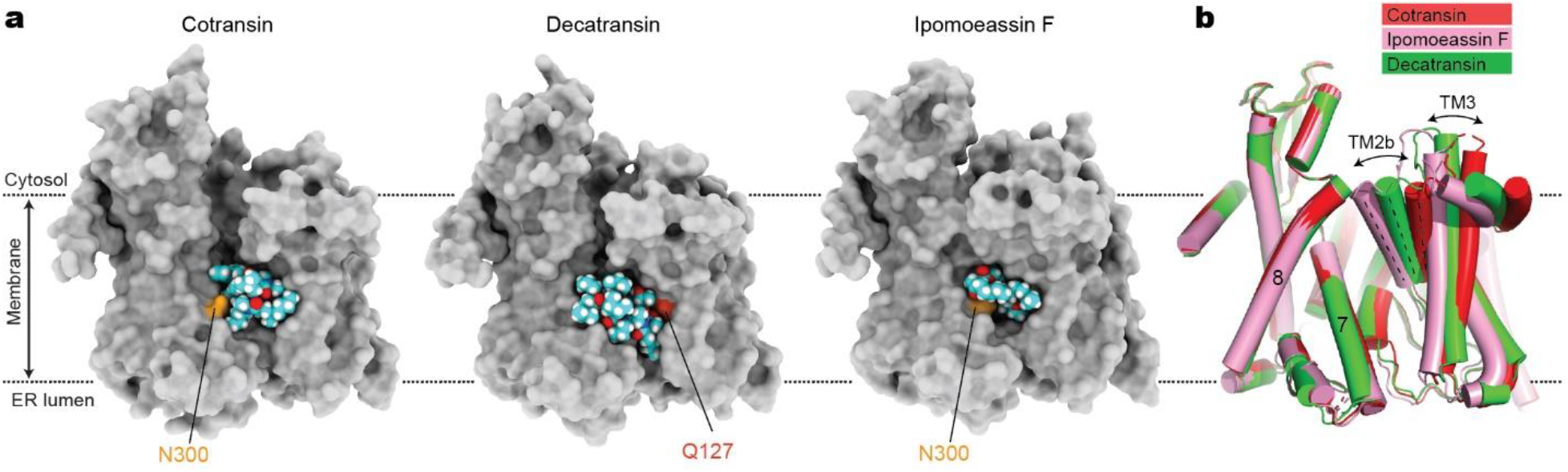
Structural plasticity of the inhibitor-binding pocket. **a**, The inhibitor-binding pocket of Sec61 and bound inhibitors are shown in surface (protein) and sphere (inhibitors) representations. Conserved polar amino acids N300 and Q127 at the inhibitor binding site (also see Fig. 3) are indicated in light and dark orange, respectively. Note that part (cinnamate moiety) of ipomoeassin is deeply buried inside the channel and invisible in this representation. **b**, Superposition of the Sec61 structures shown in **a**. Note differences in the lateral gate opening due to the varying position of the N-terminal half of Sec61α, particularly TMs 2b and 3. For other inhibitors, see Supplementary Fig. S5.

Our structures show that inhibitors bind to the lateral gate in one of these partially open states to form a tight fit with the pocket. Compared to natural inhibitors, the interfaces of CADA and ESI to Sec61 seem less extensive, possibly explaining for lower (micromolar-range) affinities of these synthetic inhibitors (Supplementary Fig. S5).

In addition to the lateral gate, the plug and pore ring critically participate in binding of all inhibitors. The partially open lateral gate of inhibited Sec61 is reminiscent of conformations observed with substrate-engaged Sec61. In fact, the inhibitor binding site largely coincides with where a signal sequence docks upon the insertion of a substrate protein into the channel^13,14,46^. However, one crucial difference exists between polypeptide substrates and inhibitors: unlike the signal sequence, all inhibitors also form a direct contact with the plug in a closed position through hydrophobic moieties (Figs. 1 and 3). Many inhibitors even further intercalate into the dilated, crescent-shaped pore ring and interact with pore-ring residues (Ile81, Val85, Ile179, Ile183, Ile292, and/or Ile449). In the cases of mycolactone and ESI, their extended chain penetrates deeply into the channel interior and occupies a substantial space of the channel’s cytosolic funnel (Fig. 3 and Supplementary Figs. S5 and S6). These parts of mycolactone and ESI are known to be critical for their inhibitory activity^23,40^.

**Figure 3.**
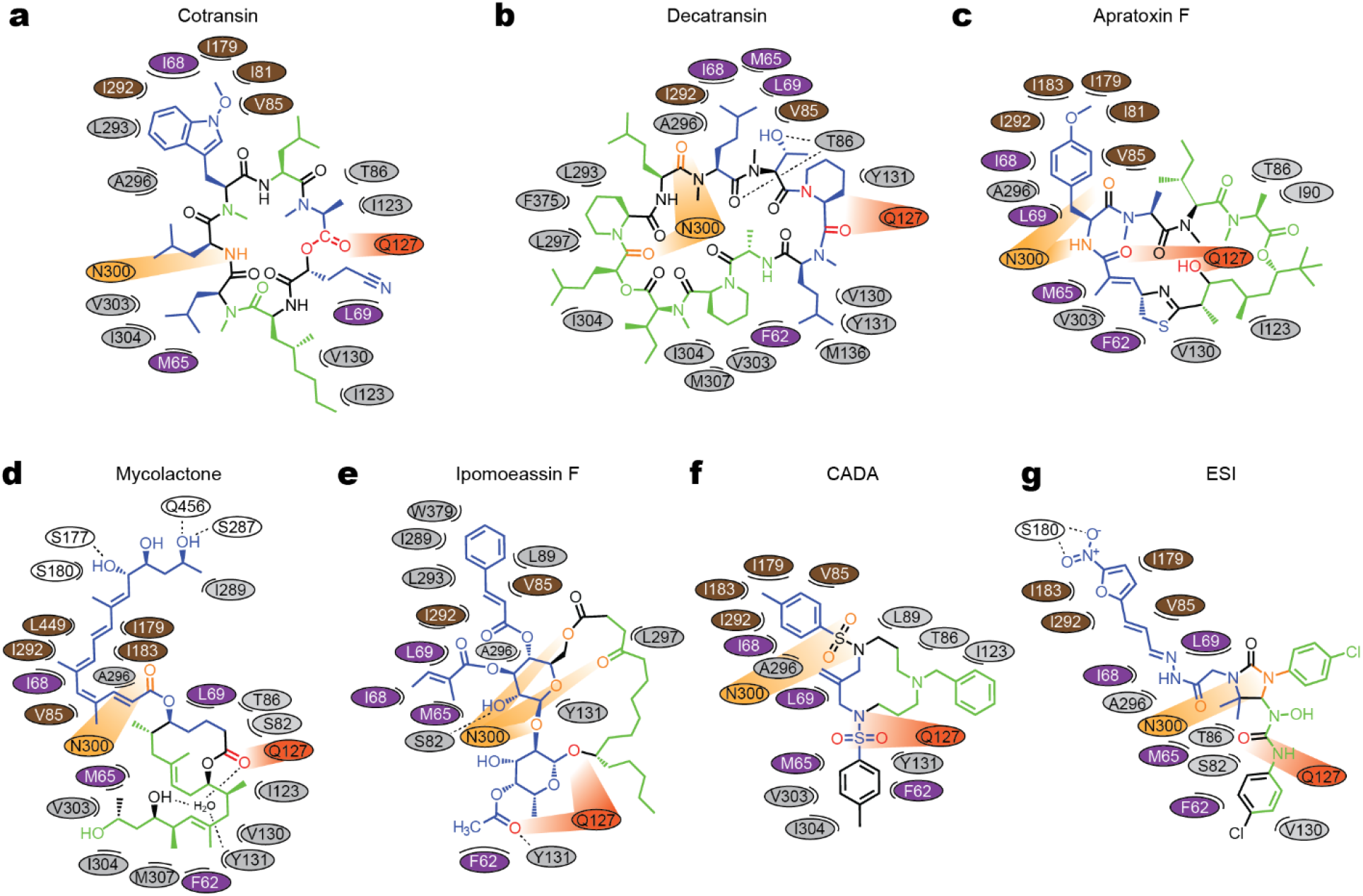
Maps for interactions between Sec61 and inhibitors. Chemical structures of inhibitors and amino acids (ovals) in the immediate vicinity are drawn in a two-dimensional representation. For actual 3D structures, see Supplementary Fig. S6. Different colors were used for ovals to indicate regions in Sec61α: purple–plug, brown–pore ring, gray–lateral gate, light and dark oranges–polar cluster Q127/N300, and white–others. In chemical diagrams of the inhibitors, main lipid-exposed parts are in green whereas channel-facing parts are in blue. Dashed lines indicate putative hydrogen bonds. Note that in the mycolactone-bound structure, a water molecule coordinated by Sec61 and mycolactone was observed in the pocket.

### Structures of Sec61 inhibitors and interactions with Sec61

Except for cotransin and apratoxin, the structures of which were determined in organic solvents by NMR spectroscopy or X-ray crystallography^47-49^, 3D structures of most Sec61 inhibitors were unknown. Our cryo-EM structures now reveal their 3D structures in association with the Sec61 channel. Notably, conformations of cotransin and apratoxin F in our cryo-EM structures are highly similar to those structures determined in organic solvent^47-49^. This might be because the inhibitor-binding site in Sec61 forms a markedly hydrophobic environment. Particularly, the pocket is open towards the lipid phase (Figs. 1 and 2), and thus, all inhibitors are expected to interact with hydrocarbon tails of membrane lipids. The lipid-exposed parts of inhibitors are predominantly hydrophobic (Fig. 3). Similarly, the parts of inhibitors that face the Sec61 channel are mostly hydrophobic as they form contacts with hydrophobic side chains from the lateral gate, plug, and pore ring of Sec61α.

While van der Waals interactions between apolar groups of inhibitors and Sec61 seem to be dominant contributors to inhibitor binding, our cryo-EM structures also show a recurring pattern of polar interactions between Sec61 and inhibitors. In the closed channel, the lateral gate contains a conserved polar cluster halfway across the membrane, formed mainly by the side chain amide groups of Gln127 (Q127) in TM3 and Asn300 (N300) in TM7. Mutations in this polar cluster has been shown to affect the energetics of channel gating^50^. In the inhibitor-bound structures, Q127 and N300 are separated by lateral gate opening, but instead they do form polar interactions with certain oxygen and nitrogen atoms in the backbones of the inhibitors. Given that these prong-like polar interactions are present in a predominantly hydrophobic milieu, it is likely that they substantially strengthen inhibitor binding at the pocket (see below).

### Mutations in Sec61 conferring resistance to inhibitors

Several point mutations in Sec61α have been found to confer resistance to Sec61 inhibitors^32,33,36-38,42,44^. These mutations are mostly located in the plug and the lateral gate. Given the direct interactions between inhibitors and these parts, disruption of the inhibitor binding surface could be a mechanism for these mutations. However, it has also been suggested that mutations may work indirectly through altering the conformation of the channel^44^. Indeed, extensive biochemical studies of the Sec61/SecY complexes have well established that mutations in the lateral gate, plug, and pore ring often change the gating behavior of the channel^50-52^. The best-known examples are *prl* mutations that give rise to relaxed client selectivity through increased propensity of channel opening. Thus, this phenotypic complexity has obscured how Sec61 mutations confer resistance to inhibitors. Moreover, positions of the identified mutations were often redundant and sparse, limiting detailed investigation of their mechanisms.

To biochemically probe inhibitor-binding sites in the Sec61 complex, we conducted a comprehensive mutational analysis fully blinded from our cryo-EM study. We focused on two inhibitors cotransin and ipomoeassin F, which were readily available to us. In addition to anti-proliferation activities on mammalian cancer cell lines, these compounds also cause growth retardation of yeast cells in a Sec61α (Sec61p)-specific manner^32^. Therefore, we tested 84 point mutations on 34 amino acid positions in yeast Sec61α for their half-maximal growth inhibitory concentration (IC_50_) (Supplementary Table S2). Positions were mainly chosen from the cytosolic funnel and lateral gate as they were likely candidates to bind inhibitors (each site was typically mutated to either Asp or Trp). This led us to identify 19 and 14 new resistance-conferring positions for cotransin and ipomoeassin F, respectively.

We then mapped the mutation positions onto the cryo-EM structures. The results clearly show that most resistance mutations are clustered around bound cotransin or ipomoeassin F (Figure 4 a and b), suggesting that their primary mechanism is through directly impairing the inhibitor-binding surface. However, some mutations (e.g., mutations equivalent to R66I/G and E78K in human Sec61α) are located at distal sites in the plug, and they may act through a conformational change in the plug domain. The plug makes a substantial contact with all tested inhibitors and is one of the most mobile parts of Sec61. Thus, altered dynamics of the plug may explain for weakened inhibitor binding.

**Figure 4.**
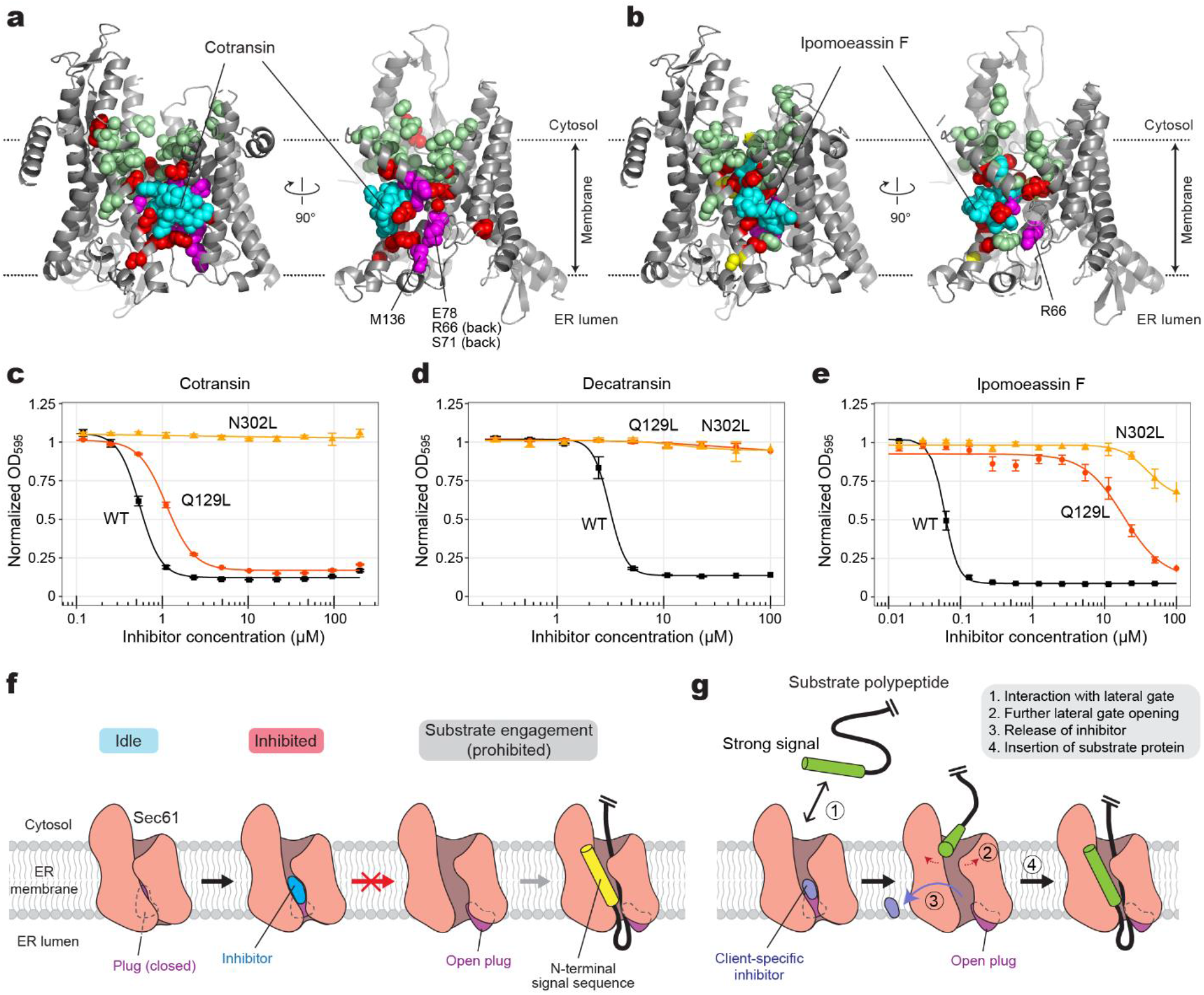
Inhibitor-resistant mutations and a model for Sec61 inhibition. **a**, Positions of mutations tested with yeast Sec61 were mapped onto the cotransin-bound structure (also see Supplementary Table S2). Left, front view; right, cutaway side view. Cotransin (cyan) and amino acid side chains are shown as spheres. Red and pale green spheres indicate positions in which mutation to Asp or Trp develops high and no cotransin resistance, respective. Magenta, positions of other resistant mutations previously reported^32,42^. **b**, as in **a**, but with ipomoeassin-F-resistant mutations. Yellow spheres additionally show positions that give rise to moderate ipomoeassin F resistance. **c**–**e**, Effects of Sec61 lateral gate polar amino acid mutations on sensitivity to cotransin, decatransin, and ipomoeassin F. Dose-response curves were generated based on the yeast growth assay (residue numbers are according to yeast Sec61). We note that the assay could not be performed for other inhibitors due to their poor response in yeast. **f**, General model for the mechanism of Sec61 inhibitors. Inhibitors bind to Sec61 in a partially open conformation and precludes the plug from opening. This prevents substrate polypeptide insertion. **g**, A proposed model for client-specific inhibition. Certain client-specific inhibitors may allow an interaction between strong signals (e.g., TM signal anchors) and the channel such that the signal sequence/anchor is wedged into the partially open lateral gate. This would further open the lateral gate to cause release of the inhibitor. Inhibitors forming less interactions with the pore and plug, rendering the lateral gate into a more open conformation, and/or displaying a weaker overall affinity are likely to be overcome by this way.

Our mutational analysis also confirms importance of polar interactions at the binding site. We found that an N302L mutation in yeast Sec61α (equivalent to N300L in human Sec61α) confers strong resistance to cotransin, decatransin, and ipomoeassin F (Fig. 4 c–e). A Q129L mutant (equivalent to Q127L in human Sec61α) showed strong to intermediate resistance to decatransin and ipomoeassin F while only little effects on cotransin. This difference may be explained in part by the structural observation that the side-chain amide of N300 more directly faces toward these inhibitors than that of Q127 (Supplementary Fig. S6).

### Mechanism of Sec61 inhibition and discussion

Our study reveals how Sec61 inhibitors interact with the channel and block the protein translocation. Remarkably, all seven tested inhibitors were found to bind to the same site in the channel formed by a partially open lateral gate and the fully closed plug domain, suggesting that this mode of interaction provides possibly the most effective mechanism for small molecules to inhibit the Sec61 channel. Among all known major Sec61 inhibitors to date, coibamide A is the only compound that was not included in the present study. However, given the previous observations that it competes with apratoxin A and mycolactone for Sec61 binding and that its resistant mutation could be found also in the plug^38^, coibamide A is likely to bind to the same or an overlapping site. We also note that the mycolactone model proposed in the previous medium-resolution cryo-EM study^44^ differs in both position and conformation from those we found in our study. Our data suggest that the density feature previously assigned as mycolactone is unlikely to be mycolactone. During the preparation of this manuscript, a medium-resolution cryo-EM structure of the mammalian Sec61 channel in association with a ribosome and a cotransin derivative has been reported^53^. While the overall structure of the channel and the location of the binding pocket seem consistent with ours, we note that the orientation of the inhibitor model is substantially different from that of cotransin from our study. This discrepancy is more likely due to a limited map resolution of the ribosome-Sec61 structure, although we cannot rule out a possibility that it may originate from minor structural differences between the two compounds.

Despite their distinct chemical structures, some common features among the inhibitors could be derived from our results. First, the inhibitors have two major clusters of hydrophobic moieties, one arranged to interact with the plug and the lateral gate, and the other with membrane lipids. The Sec61-facing sides are characterized by strong surface complementarity for the binding pocket, while the lipophilicity of the other parts would also contribute to efficient binding as the pocket exists within the plane of the membrane. Second, all inhibitors form polar interactions between their backbone and the side chains of the lateral gate (Asn300, Gln127, and/or Thr86 of Sec61α). We found that this is crucial for Sec61 binding affinity. These polar groups of inhibitors would also provide some water solubility of the compounds. Third, certain inhibitors, such as mycolactone and ESI, further penetrate the cytosolic funnel of Sec61 forming additional polar and hydrophobic interactions therein. These interactions likely contribute to more stable binding and broad-spectrum inhibition.

Our data indicate that all known Sec61 inhibitors block the protein translocation process commonly by locking both lateral and lumenal gates of Sec61 into translocation-incompetent conformations (Fig. 4f). Although the lateral gate stays partially open, it does not provide a sufficient space for a signal sequence/anchor to pass. Importantly, the lumenal gate, i.e., the plug, remains fully closed such that the client polypeptide cannot insert into the pore. Overall, all three key gating elements—the lateral gate, plug, and pore ring—are cemented together by the inhibitor at their interface, thereby prohibiting their concerted opening required for the client protein insertion.

Although further investigations would be necessary, our comparative analysis also hints at why certain inhibitors exhibit client-dependent translocation inhibition. Cotransin and CADA have been shown to be less effective in blocking translocation of client proteins containing a stronger targeting signal, such as a signal sequence with higher hydrophobicity or a TM signal anchor^54-58^. Our structures show that in the cotransin-bound structure, the lateral gate adopts a relatively more open conformation on the cytosolic side (Fig. 2). This may allow certain hydrophobic interactions between the lateral gate and the incoming signal sequence/anchor (Fig. 4g). A stronger interaction exerted by a stronger targeting signal probably tends to further pry open the lateral gate, promoting the inhibitor to be released. Although the lateral gate of the CADA-bound structure is not as wide as that of cotransin-bound Sec61, its relatively low binding affinity (∼0.2 μM) might facilitate certain hydrophobic signals to overcome inhibition. On the other hand, those inhibitors that deeply insert into the pore and cytosolic funnel of the channel may tend to exert broad-spectrum inhibition by additionally impeding client insertion into the pore. Mycolactone and ESI fall into such a category.

It remains unclear whether binding of an inhibitor requires prior opening of the Sec61 channel. In our chimeric complex, the lateral gate is partially opened by Sec63. In co-translational translocation, it has been generally thought that the ribosome docking alone does not open the lateral gate to a considerable extent^11^, which seems necessary for inhibitor binding. However, a transient breathing motion of the channel might allow inhibitors to bind. Single-molecule fluorescence studies of the bacterial SecY channel have indicated that the lateral gate spontaneously fluctuates between closed and open states without any binding partner^59,60^. Thus, it is possible that inhibitor binding may not require priming or partial opening of the channel induced allosterically by the ribosome or Sec63.

Lastly, the rich structural and mechanistic knowledge we provide here can facilitate structure-guided design of Sec61 inhibitors. The Sec61 channel has been considered as a promising target for therapeutic intervention due to its essential role in production of many cytokines, surface receptors, and cell adhesion molecules that are clinically relevant. Nevertheless, currently available Sec61 inhibitors would need further structural optimizations to improve their effectiveness and pharmacological properties while reducing undesired cytotoxicity. Our new approach enabling high-resolution structural analysis of human Sec61 and bound ligands would accelerate efforts to understand the mechanisms of new Sec61 inhibitors and optimize previously identified molecules.

## Acknowledgments

We thank Dan Toso for support for electron microscope operation, Guanghui Zong for ipomoeassin F synthesis, Philippe Mathys and Ralph Riedl for help acquiring the IC50 data. E.P. was supported by the Vallee Scholars Program (E.P.) and Pew Biomedical Scholars Program. S.I. and L.W. were supported by a National Institutes of Health training grant (5T32GM008295). M.S and T.J. were supported by the Swiss National Science Foundation (31003A-182519). N.B was supported by Fondation Raoul Follereau and Fondation Pour Le Développement De La Chimie Des Substances Naturelles Et Ses Applications. W.Q.S. was supported by an AREA grant from National Institutes of Health (GM116032). C.F. and L.X. were supported by the Ohio State University.

## Author contributions

E.P. conceived the project and supervised the cryo-EM study. L.W. and S.I. cloned the chimeric Sec construct and prepared protein samples. S.I., L.W., and E.P. collected and analyzed cryo-EM data and built atomic models. R.S. helped purification of the human Sec complex and cloning of the chimeric Sec complex. T.J., M.S., and D.H. performed the yeast mutational study. D.H. provided cotransin and decatransin. C.F. and L.X. provided apratoxin F. W.S. provided ipomoeassin F. N.B. provided mycolactone. All authored contributed to interpret results. E.P. wrote the manuscript with input from all authors.

## Competing interests

The remaining authors declare no competing interests.

## Legends for Supplementary Figures

**Supplementary Figure S1. Cryo-EM analysis of the yeast and human Sec complexes**.

**a**, A schematic of the single-particle cryo-EM analysis of the yeast Sec (*Sc*Sec) complex incubated with cotransin. Note that the particles were sorted into two 3D classes, with and without Sec62, due to partial occupancy of Sec62. **b**, 3D reconstructions of the *Sc*Sec complex with and without *Sc*Sec62 (shown in yellow). No cotransin-like density was observed in either class. For this experiment, we used a pore ring mutant (PM; M90L/T185I/M294I/M450L) that stabilize the plug towards a closed conformation. **c**, Purification of the human Sec (*Hs*Sec) complex. Shown is a Superose 6 size-exclusion chromatography elution profile with fractions analyzed on a Coomassie-stained SDS gel. Note that under the used purification condition, *Hs*Sec62 does not co-purify at a stoichiometric ratio or stably comigrate with the Sec61–Sec63 complex. The fractions indicated by gray shade were used for cryo-EM. MW standards: Tg, thyroglobulin; F, ferritin; Ald, aldolase. **d**, A schematic of the single-particle analysis of *Hs*Sec complex incubated with cotransin. Due to a poor refinement result from nonuniform refinement in cryoSPARC, the final reconstruction was obtained by the ab-initio refinement function of cryoSPARC (see **f**). **e**, Representative 2D classes of the *Hs*Sec complex. Diffuse cytosolic features of Sec63 (green arrowheads) suggest its flexibility or disorderedness. **f**, The 3D reconstruction of the *Hs*Sec complex. A putative cotransin feature (cyan) is visible at the lateral gate.

**Supplementary Figure S2. Cryo-EM analysis of the chimeric Sec complex in an apo form**.

**a**, Purification of the chimeric Sec complex reconstituted in a peptidisc. Left, Superose 6 elution profile; right, Coomassie-stained SDS gel of the peak fraction. The fraction marked by gray shade was used for cryo-EM. Asterisks, putative species of glycosylated *Sc*Sec71. **b**, A schematic of the cryo-EM analysis of the chimeric Sec complex in an apo state. **c** and **d**, Distributions of particle view orientations in the final reconstructions of Classes 1 (c) and 2 (d). **e** and **f**, Fourier shell correlation (FSC) curves and local resolution maps of the final reconstructions. **g**, Superimposition of the Class 1 and 2 atomic models (based on the cytosolic domains) shows a slight difference in relative positions between Sec63-Sec71-Sec72 and the Sec61 complex. **h**, Side views showing the contact between the engineered cytosolic loops of Sec61α and the FN3 domain of *Sc*Sec63. Note that in Apo Class 2, the contact is more poorly packed than Class 1.

**Supplementary Figure S3. Cryo-EM analysis of the chimeric Sec complex in an inhibitor (apratoxin F)-bound form**.

**a**, Images of a representative micrograph and particles of the apratoxin F-bound chimeric Sec complex. Scale bar, 10 nm. **b**, A schematics of the cryo-EM analysis of the apratoxin F-bound chimeric Sec complex. **c**, Representative 2D classes of the apratoxin F-bound Sec complex. **d**, Distribution of particle view orientations in the final reconstruction. **e**, The FSC curve and local resolution map of the final reconstruction (full Sec complex map). **f**, As in e, but for the map from focused (local) refinement. **g**, Segmented density maps of the apratoxin F-bound Sec61α subunit. **h**, Segmented density features of bound natural inhibitors.

**Supplementary Figure S4. FSC curve and local resolution maps of inhibitor-bound Sec complexes**.

As in Supplementary Figure S3 e and f, but for all other inhibitor-bound structures.

**Supplementary Figure S5. Variation in the extent of lateral gate opening in inhibitor-bound structures**.

As in Fig. 2 a and b, but showing other inhibitor-bound structures. In all panels showing a lateral gate comparison, cylindrical representations in red and pink are the cotransin- and ipomoeassin F-bound structures, respectively, whereas the representation in green is the structure with the indicated inhibitor.

**Supplementary Figure S6. 3D maps for interactions between Sec61 and inhibitors**.

Shown are stereo-views into the inhibitor-binding site. Inhibitors and adjacent protein side chains are shown in a stick representation together with Cα traces for TM2b, TM3, TM7, and the plug. The views are roughly similar between the different structures but adjusted for each structure for more clear representations. The following colors are used to differentiate parts: brown, pore ring residues; magenta, plug; lighter orange; N300, darker orange, Q127. All inhibitors are shown in cyan with certain atom-dependent coloring (nitrogen-blue, oxygen-red, sulfur-yellow, and chlorine-green).

## Materials and Methods

### Sec61 Inhibitors

Isolation of cotransin (previously referred to as “Compound 2”) and decatransin from fugal species have been described previously^32^. For apratoxin F, ipomoeassin F, and mycolactone, we used synthetic versions. Synthesis of apratoxin F (ref. ^61,62^), ipomoeassin F (ref. ^63,64^), mycolactone (ref. ^65^) has been as described previously. We note that apratoxin F and its more commonly studied analog apratoxin A have only a minor structural difference and both are known to exhibit comparable IC_50_ values on mammalian cancer cell lines^62^. We also note that the used synthetic mycolactone is a 4:1 mixture of two epimers at C12 in favor of the natural configuration. CADA and ESI were purchased from Calbiochem. Inhibitors were dissolved in dimethyl sulfoxide (DMSO) at 10 mM (for decatransin, ipomoeassin F, and mycolactone), 20 mM (for cotransin, apratoxin F, and CADA), or 50 mM (for ESI) before use.

### Plasmid constructs for cryo-EM studies

The plasmids and yeast strain to express the *S. cerevisiae* Sec complex have been described previously^15,17^. To express the human Sec complex in *Spodoptera frugiperda* (Sf9) cells, we modified a Bac-to-Bac baculovirus expression vector (Invitrogen) adapting the multigene-expression approach from MoClo Yeast ToolKit (YTK)^66^ as follows. First, we generated part plasmids for a baculovirus polyhedrin (PH) promoter and a SV40 polyA signal, and an accepter plasmid (pBTK1) consisting of the backbone of pFastBac-1 (including a Tn7L element, an ampicillin resistance gene, a pUC *E. coli* origin of replication, a Tn7R element and a gentamycin resistance gene) and a *Bsm*BI–superfolder GFP (sfGFP)–*Bsm*BI acceptor cassette from pYTK096 (ref. ^66^). Gene fragments encoding human Sec subunits were chemically synthesized and individually cloned into the entry plasmid pYTK001 as coding sequence (CDS) parts. Amino acids sequences of human (denoted by “*Hs*”) Sec61, Sec62, and Sec63 subunits are from the following entries in UniProt: P61619 (S61A1_HUMAN) for *Hs*Sec61α, P60468 (SC61B_HUMAN) for *Hs*Sec61β, P60059 (SC61G_HUMAN) for *Hs*Sec61γ, Q99442 (SEC62_HUMAN) for *Hs*Sec62, and Q9UGP8 (SEC63_HUMAN) for *Hs*Sec63. For the pYTK001-*Hs*Sec63 plasmid, a DNA segment encoding a human rhinovirus (HRV) 3C-cleavable linker (amino acid sequence: GAGSNS*LEVLFQGP*TAAAA; italic, HRV 3C cleavage site) and an enhanced green fluorescence protein (eGFP) were inserted immediately before the stop codon of *Hs*Sec63. To generate single Sec gene expression cassettes, each Sec subunit CDS was assembled with connectors (from pYTK003–007 and pYTK067–072), the PH promoter, and the SV40 terminator into pYTK095 using *Bsa*I Golden Gate cloning. Then, all Sec subunit expression cassettes were assembled into pBTK1 using *Bsm*BI Golden Gate cloning. In this multigene plasmid, the expression cassettes were arranged in the following order: PH-*Hs*Sec61α-SV40 | PH-*Hs*Sec61γ-SV40 | PH-*Hs*Sec61β-SV40 | PH-*Hs*Sec63-3C-eGFP-SV40 | PH-*Hs*Sec62-SV40.

The plasmid expressing the human-yeast chimeric Sec complex were made similarly to the human Sec complex plasmid with modifications of pYTK95 *Hs*Sec61α and *Hs*Sec63 expression constructs as follows. To modify Sec61α, two substitution mutations were introduced in cytosolic loops of *Hs*Sec61α using PCR to replace (1) amino acid residues 263–278 (VDLPIKSARYRGQYNT) with the corresponding yeast sequence (residues 265–280; YELPIRSTKVRGQIGI) and (2) amino acid residues 394–411 (LKEQQMVMRGHRETSMVH) with amino acids 395–412 of *Sc*Sec61 (FKDQGMVINGKRETSIYR; “*Sc*” denotes *Saccharomyces cerevisiae*). The substitutions in *Hs*Sec63 were introduced using Gibson assembly by first substituting amino acid residues 30–96 (ATY…VKK) with amino acids 29–93 of *Sc*Sec63 (MTL…RRN), followed by substitution of residues 215 to the C-terminus (SIR…stop) with the corresponding sequence from *Sc*Sec63 (residues 246–stop; TQS…stop). Fragments of *Sc*Sec63 were amplified from genomic DNA of yeast strain BY4741. In the multigene pBTK1 construct of the chimeric Sec complex, *Hs*Sec62 cassette was omitted, and instead, the cassettes for *Sc*Sec71 (PH-ScSec71-SV40) and *Sc*Sec72 (PH-ScSec72-SV40) were added. The CDS fragments of *Sc*Sec71 and *Sc*Sec72 were amplified by PCR from genomic DNA of yeast strain BY4741 and cloned into pYTK001. Like other single subunit expression plasmids, *Sc*Sec71 and *Sc*Sec72 CDSs were assembled into pYTK095 together with the PH promoter and the SV40 polyA signal before use for the *Bsm*BI assembly.

### Protein Expression

Baculovirus bacmids encoding the human or chimeric Sec complex were generated by transforming the respective pBTK1 plasmid into the DH10Bac *E. coli* cells (Invitrogen) according to the manufacturer’s instructions. Bacmids were isolated using a DNA midiprep kit (Epoch Life Science). 40 mL of a Sf9 suspension culture were prepared in ESF921 medium (Expression Systems) to a density of ∼1.5 M/mL. 40 μg bacmid DNA were mixed with 80 μg PEI Max transfection reagent (PolySciences) in 4 mL Dulbecco’s phosphate-buffered saline (DPBS). After incubating at 22°C for 20–30 minutes, the DNA:PEI mixture was added to the culture. Supernatant containing P1 virus was harvested ∼4 days post transfection and stored at 4°C after supplementing 5% FBS (Gibco). Expression of the Sec complex was carried out by adding 0.5 mL P1 virus to 0.7 L of Sf9 cells at density of ∼1.5 M/ml that were prepared in a 2-L baffled flask. Cells were harvested by centrifugation typically two to three days post-infection upon verifying uniform expression of green fluorescence under microscope. Cell pellets were frozen in liquid nitrogen and stored at -80°C until use.

### Protein Purification

The yeast Sec complex was purified from yeast strain ySI8 (ref. ^17^). This strain expresses a “pore mutant (PM)” version of *Sc*Sec61, the pore ring residues of which were mutated to amino acids corresponding to *Hs*Sec61α (M90L/T185I/M294I/M450L). The yeast Sec complex was purified as described previously^15,17^. After Superose 6 (GE Life Sciences) size-exclusion chromatography, the purified protein was concentrated to ∼4 mg/mL in 20 mM Tris pH 7.5, 100 mM NaCl, 1mM EDTA, 2 mM DTT, and 0.02% glycol-diosgenin (GDN; Anatrace) and mixed with 100 μM cotransin for 0.5–1 h before preparing cryo-EM grids.

To purify the human Sec complexes, Sf9 cell pellets were first resuspended in lysis buffer containing 50 mM Tris-HCl pH 7.5, 200 mM NaCl, 2 mM dithiothreitol (DTT), 1 mM ethylenediaminetetraacetic acid (ETDA) supplemented with protease inhibitors (5 μg/ml aprotinin, 5 μg/ml leupeptin, 1 μg/ml pepstatin A, and 1.2 mM PMSF). All subsequent steps were carried out in ice or at 4°C. The cells were broken with a glass Dounce homogenizer using ∼100 strokes. After removing large debris by brief centrifugation (4,000 g, 10 min), membranes were pelleted by ultracentrifugation for 1.5 h (125,000 g, Beckman Type 45 Ti). The membrane pellet was resuspended in ∼10 pellet volumes of lysis buffer supplemented with 5 μM cotransin. Membranes were solubilized by an addition of 1% lauryl maltose neopentyl glycol (LMNG; Anatrace) and 0.2% cholesteryl hemisuccinate (CHS; Anatrace) for 2 h. Then, the lysate was clarified by ultracentrifugation at 125,000 g for 1 h. The clarified lysate was then supplemented with 2 μg *Serratia marcescens* nuclease and incubated with home-made anti-GFP nanobody Sepharose beads for 1.5 h. Beads were washed with wash buffer containing 25 mM Tris-HCl pH 7.5, 100 mM NaCl, 2 mM DTT, 1 mM ETDA, 0.02% GDN, and 5 μM cotransin (hereafter, 5 μM cotransin was included in all buffers). The complex was eluted by incubating beads with the HRV 3C protease overnight. The eluate was collected and concentrated to ∼10 mg/ml by Amicon Ultra (cutoff 100 kDa). The sample was then injected to a Superose 6 increase column (GE Life Sciences) equilibrated with the wash buffer. Peak fractions were pooled and concentrated to ∼6 mg/mg, before preparing cryo-EM grids.

The chimeric Sec complex was purified similarly using the method to purify the human Sec complex but with minor modifications. First, the Sec complex was purified without supplementing Sec61 inhibitors during purification (inhibitors were added to the purified Sec complex before cryo-EM grid preparation). Second, to solubilize membranes, 1% n-dodecyl-β-D-maltopyranoside (DDM; Anatrace) and 0.2% CHS was used instead of LMNG/CHS. For column wash, the buffer contained 0.02% DDM and 0.004% CHS instead of GDN. Third, the Sec complex was reconstituted into a peptidisc^45^ as follows. After concentrating the eluate from GFP-nanobody beads to ∼10 mg/ml, the Sec complex was mixed with the peptidisc protein (Peptidisc Lab) at a weight ratio of 1.5:1 (peptidisc to Sec). After incubating for 1 h, the mixture was injected into a Superose 6 Increase column equilibrated with 25 mM Tris-HCl pH 7.5, 100 mM NaCl, 2 mM DTT and 1 mM ETDA. Peak factions were pooled and concentrated to ∼10 mg/ml (∼52 μM), and one of the Sec61 inhibitors was added for ∼1 h before preparing cryo-EM grids. The inhibitor concentrations used were: 100 μM for cotransin, 100 μM for decatransin, 100 μM for apratoxin F, 100 μM for ipomoeassin F, 100 μM for mycolactone, 200 μM for CADA, and 500 μM for eeyarestatin I.

### Cryo-EM data acquisition

Immediately prior to preparing cryo-EM grids, 3 μM Fos-Choline-8 (Anatrace) was added to the protein sample. The sample was then applied to a gold Quantifoil R 1.2/1.3 holey carbon grid (Quantifoil) that was glow discharged for 35 sec using PELCO easiGlow glow discharge cleaner. The grid was blotted for 3–4 sec using Whatman No. 1 filter paper and plunge frozen using Vitrobot Mark IV (FEI) set at 4°C and 100% humidity.

The yeast Sec complex dataset (1,578 movies) was acquired on FEI Talos Arctica electron microscope operated at an acceleration voltage of 200 kV, with Gatan K2 Summit direct electron detector. A magnification of 36,000x under super resolution mode (with the physical pixel size of 1.14 Å) was used with a nominal defocus range that was set between -0.8 to -2.2 μm. Each micrograph was composed of 42 frames with total exposure of 50 e^-^/pixel.

The human Sec complex dataset (3,499 movies) was collected on FEI Titan Krios G2 microscope operating at an acceleration voltage of 300 kV and equipped with a Gatan Quantum Image Filter (slit width of 20 eV) and a Gatan K3 direct electron detector. A magnification of 64,000x under the super-resolution mode (with physical pixel size of 0.91 Å) was used at a defocus range that was set between -0.8 and -2.0. Each micrograph was composed of 42 frames with total exposure of 50 e^-^/pixel. Exposures were performed with beam shifts onto 9 holes (3 by 3) per stage movement.

All chimeric Sec complex datasets were acquired on an FEI Titan Krios G3i electron microscope operating at an acceleration of 300 eV, with a Gatan K3 Summit direct electron detector and a Gatan Quantum Image Filter (with 20 eV slit width). A magnification of 81,000x under the super-resolution mode (with physical pixel size of 1.05 Å) was used at a defocus range that was set between -0.8 and -2.0. Each micrograph was composed of 50 frames with a total exposure of 50 e^-^/pixel. Exposures were performed with beam shifts onto 9 holes (3 by 3) per stage movement (often acquiring movies for two non-overlapping areas per hole). All datasets were acquired using SerialEM software^67^.

### Cryo-EM image analysis

Preprocessing of the movies and particle image extraction were done using Warp^68^. Motion correction and CTF estimation were performed on images divided to 7 × 5 tiles, and particles (256 × 256 pixels) were picked by the BoxNet algorithm in Warp. All subsequent image processing procedures were perform using cryoSPARC v3.3 (ref. ^69^).

Cotransin treated pore mutant *Sc*Sec complex: A data processing flowchart diagram is shown in Supplementary Fig. S1a. A dataset of 528,128 auto-picked particles was classified into fifty 2-D class averages. Using visual inspection of the output, classes that represented empty micelles or poor-quality classes were removed and particles grouped into well resolved classes corresponding to a single copy of the full Sec complex were selected (385,686 particles). Three ab-initio 3-D maps were then generated in cryoSPARC using the selected particles, followed by heterogeneous refinement. One 3-D class with 274,913 particles refined to a density map exhibiting defined Sec complex features. Non-uniform refinement of the particles in this class yielded a consensus map with 3.9-Å overall resolution. The particles were further separated into two 3-D classes using a heterogeneous refinement with inputs of the consensus map and the consensus map with manually erased Sec62. After a subsequent round of non-uniform refinement 174,058 particles yielded a map of *Sc*ScSec complex with Sec62 at 4.0-Å overall resolution, and 100,855 particles yielded a map of *Sc*Sec complex without Sec62 at 4.2-Å overall resolution.

Cotransin-bound wildtype *Hs*Sec complex: A data processing flowchart diagram is shown in Supplementary Fig. S1d. A dataset of 601,465 auto-picked particles was classified into seventy 2-D class averages. Selected classes yielded 330,005 particles that were then reconstructed into three 3-D classes using ab-initio reconstruction followed by heterogeneous refinement. One major class, with 202,946 particles, was selected for further refinement. Non-uniform refinement of this class resulted in a reconstruction only at 7.4-Å resolution due to poor image alignment. Thus, for the final map, we used the ab-initio reconstruction method (without splitting the particle sets for half maps) with the maximal refinement resolution manually set to 5.0-Å (Supplementary Fig. S1f).

Apo chimeric Sec complex: A data processing flowchart diagram is shown in Supplementary Fig. S2b. Using 2-D classifications starting with 616,121 auto-picked particles, we selected 363,027 particles for 3-D reconstruction. Following an ab-initio refinement step generating four initial maps and a heterogeneous refinement step we identified two major 3-D classes with distinguishable full Sec complex features. Each of these classes were refined using non-uniform refinement, local CTF refinement, and another round of non-uniform refinement, yielding full maps of the chimeric Sec complex at overall resolutions of 2.7 and 2.8 Å from 188,637 particles (Class 1) and 147,081 particles (Class 2), respectively. The Sec61 channel was further refined by masking out the cytosolic domains of the complex and performing local refinement, yielding overall channel resolutions of 3.0 Å (Class 1) and 3.4 Å (Class 2).

Apratoxin F-bound chimeric Sec complex: A data processing flowchart diagram is shown in Supplementary Fig. S3b. Using 2-D classifications starting with 910,463 auto-picked particles, we selected 534,411 particles for 3-D reconstruction. Following an ab-initio refinement step generating four initial maps and a heterogeneous refinement step we identified two structurally indistinguishable major 3-D classes with defined full Sec complex features. The particles from the two classes were combined and refined using non-uniform refinement, local CTF refinement, and another round of non-uniform refinement, yielding a full map of the apratoxin F bound chimeric Sec complex at an overall resolution of 2.5 Å from 497,555 particles. The Sec61 channel was further refined by masking out the cytosolic domains of the complex and performing local refinement, producing an overall channel resolution of 2.6 Å.

All other inhibitor-bound datasets were processed using a workflow described for Apratoxin F-bound structure with minor variations in the numbers of classes in 2-D and 3-D classification procedures. For details, see Supplementary Figs. S7 and S8. Statistics for final refined maps are shown in Supplementary Fig. S4 and Supplementary Table S1.

### Model building and refinement

Atomic model building and refinement were performed using Coot^70^ and Phenix^71^. An initial model was built by docking an *Sc*Sec complex model (PDB ID 7KAH; ref. ^17^) into the cryo-EM map of the cotransin-bound complex using UCSF chimera^72^ and rebuilding the polypeptide chains. For building and refining of Sec61 and inhibitor models, we used maps from focused (local) refinements as they typically showed better protein side-chain and inhibitor features than full maps. The initial model was further improved by using our highest-resolution map, which was obtained from the apratoxin F-bound complex. This model was then used to build atomic models for apo and other inhibitor-bound complexes by docking the model to the map using UCSF chimera and locally adjusting it into the map in Coot. The restraint models of inhibitors were generated from SMILES strings of inhibitors using the Grade web server (http://grade.globalphasing.org) or the eLBOW tool of Phenix. The atomic models of inhibitors were then fitted into the cryo-EM map in Coot. We note that stereochemistry of decatransin has not been experimentally determined. We assumed that all amino acid residues of decatransin are in an L or S configuration based on an observation that no epimerase was found in the biosynthetic gene cluster of decatransin. The configuration of Cα of the homoleucine-derived 2-hydroxy carboxylic acid remains ambiguous^32^, but we also assumed that it is in an S configuration. The resulting model could be fitted well into the cryo-EM map. The atomic models were refined with Phenix real-space refinement using maps that were sharpened with B-factors estimated based on the Guinier plots and low-pass-filtered at their overall resolution. The refinement resolution was also limited to the overall resolution of the maps in Phenix. Structural validation was performed using MolProbity^73^. UCSF Chimera, ChimeraX (ref. ^74^), and PyMOL (Schrödinger) were used to prepare figures in the paper.

### Mutagenesis of yeast Sec61α and IC_**50**_ **measurements**

Except for the experiment shown in Fig. 4c, cotransin IC_50_ measurements were based on the yeast strain RSY1293 (matα, *ura3-1, leu2-3,-112, his3-11,-15, trp1-1, ade2-1, can1-100, sec61::HIS3*, [pDQ1]) (ref. ^75^). In strain RSY1293URA, pDQ1, i.e., YCplac111 (*LEU2 CEN*) containing the gene of an N-terminally His-tagged, otherwise wild-type Sec61α (Sec61p) with its own promoter, was exchanged for YCplac33 (*URA3 CEN*) containing the same insert. Mutations in *sec61* were introduced in pDQ1 by PCR and transformed into RSY1293URA, followed by elimination of the *URA3* plasmid containing wild-type using 5-fluoro-orotic acid. Finally, the presence of the mutation was confirmed by sequencing.

For experiments shown in Fig. 4 c–e and ipomoeassin IC_50_ measurements, we used the yeast strain BY4743Δ8a (*mat a, ura3Δ0, leu2Δ0, his3Δ1, lys2Δ0, snq2::KanMX4; pdr3::KanMX4; pdr5::KanMX4; pdr1::NAT1; yap1::NAT1; pdr2Δ; yrm1Δ; yor1Δ*) lacking eight genes involved in drug resistance (efflux pumps *SNQ2, PDR5*, and *YOR1*, and transcription factors *PDR1, PDR2, PDR3, YAP1*, and *YRM1*)^76^. This strain showed higher sensitivity to ipomoeassin F compared to RSY1293. YCplac33 containing (untagged) *SEC61* with 200 bp of its own upstream and 205 bp of its own downstream sequence was transformed into BY4743Δ8a. Genomic *SEC61* together with 194 bp 5’- and 204 bp 3’-noncoding sequence was replaced with a hygromycin resistance cassette using pAG32 (*HphMX4*) (ref. ^77^) resulting in the strain BY4743Δ9aURA (*sec61::HphMX4* [pSEC61-YCplac33]). Finally, pDQ1 containing the mutated *sec61* versions were transformed and the wild-type *SEC61* URA3 plasmid counterselected. Plasmid exchange was validated by PCR. The IC_50_ measurements for cotransin in the BY4743Δ9aURA background paralleled those in the RSY1293.

IC_50_ measurements were performed as described previously^32^ by testing log-phase cultures in 96-well microtiter plates in YPD medium with serial dilutions of the compound. The assay volume was 120 µl/well, start OD_600_ was 0.05, DMSO was normalized to 2%. Curves were calculated by taking the 19 h OD_600_ measurements and applying a log regression curve fit in TIBCO Spotfire v3.2.1.

## Notes

### Competing Interest Statement

The authors have declared no competing interest.

