## Supplementary Figures S1-S8 and Tables S1-S2 for "A common mechanism of Sec61 translocon inhibition by small molecules"

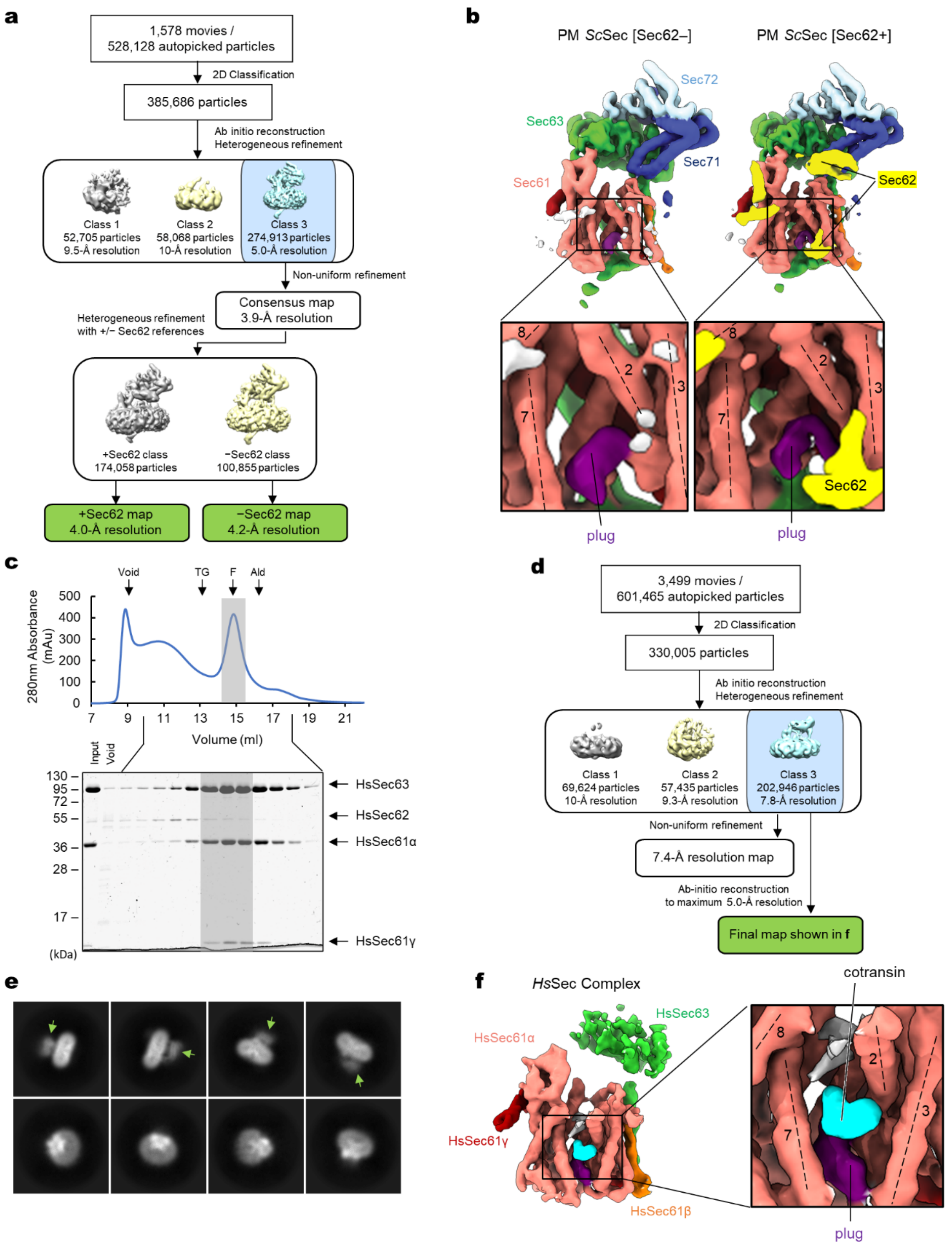


**Supplementary Figure S1. Cryo-EM analysis of the yeast and human Sec complexes**.
**a**, A schematic of the single-particle cryo-EM analysis of the yeast Sec (*Sc*Sec) complex incubated with cotransin. Note that the particles were sorted into two 3D classes, with and without Sec62, due to partial occupancy of Sec62. **b**, 3D reconstructions of the *Sc*Sec complex with and without *Sc*Sec62 (shown in yellow). No cotransin-like density was observed in either class. For this experiment, we used a pore ring mutant (PM; M90L/T185I/M294I/M450L) that stabilize the plug towards a closed conformation (ref. 17). **c**, Purification of the human Sec (*Hs*Sec) complex. Shown is a Superose 6 size-exclusion chromatography elution profile with fractions analyzed on a Coomassie-stained SDS gel. Note that under the used purification condition, *Hs*Sec62 does not co-purify at a stoichiometric ratio or stably comigrate with the Sec61–Sec63 complex. The fractions indicated by gray shade were used for cryo-EM. MW standards: Tg, thyroglobulin; F, ferritin; Ald, aldolase. **d**, A schematic of the single-particle analysis of *Hs*Sec complex incubated with cotransin. Due to a poor refinement result from nonuniform refinement in cryoSPARC, the final reconstruction was obtained by the ab-initio refinement function of cryoSPARC (see **f**). **e**, Representative 2D classes of the *Hs*Sec complex. Diffuse cytosolic features of Sec63 (green arrowheads) suggest its flexibility or disorderedness. **f**, The 3D reconstruction of the *Hs*Sec complex. A putative cotransin feature (cyan) is visible at the lateral gate.

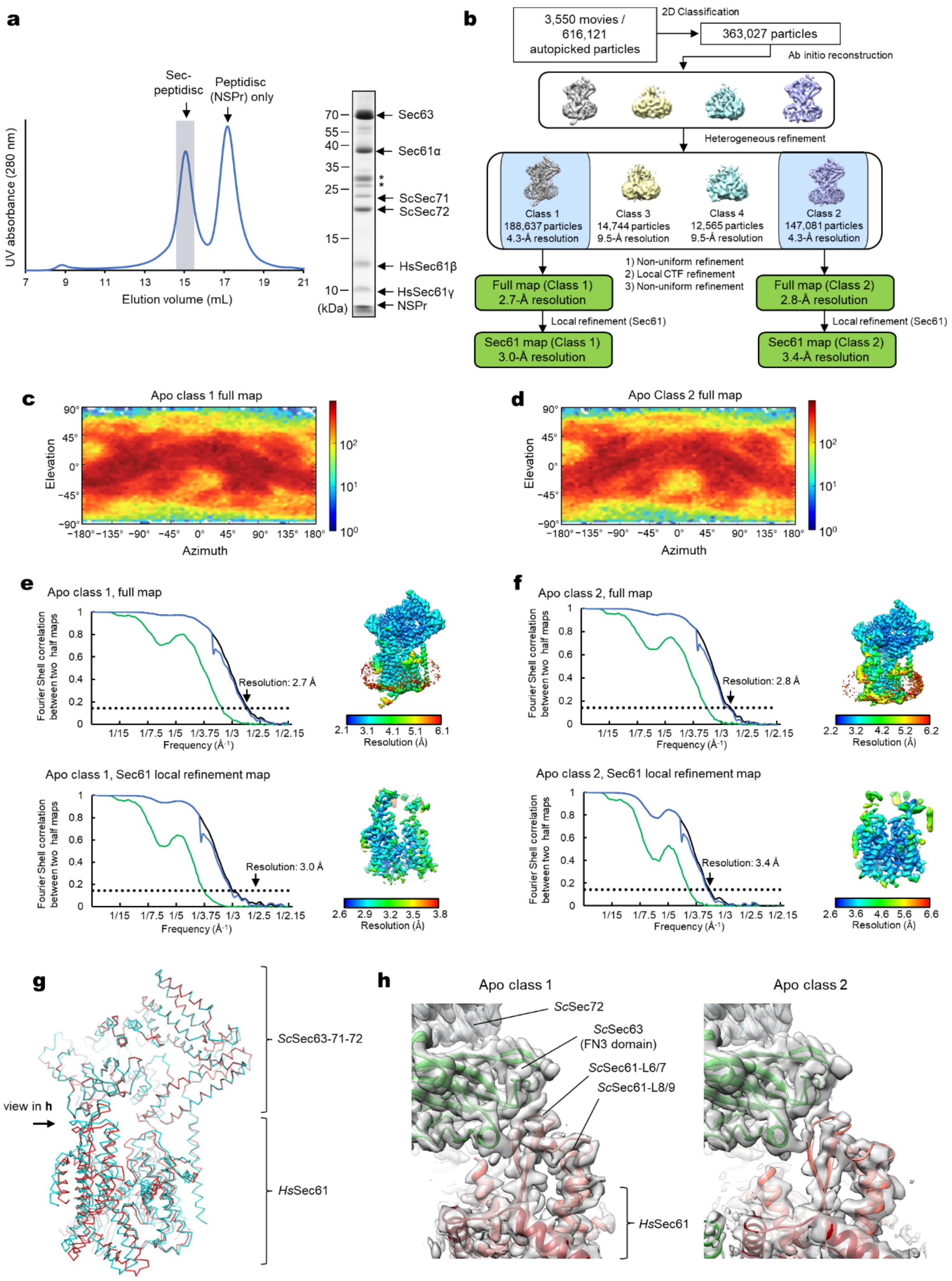


**Supplementary Figure S2. Cryo-EM analysis of the chimeric Sec complex in an apo form.**
**a**, Purification of the chimeric Sec complex reconstituted in a peptidisc. Left, Superose 6 elution profile; right, Coomassie-stained SDS gel of the peak fraction. The fraction marked by gray shade was used for cryo-EM. Asterisks, putative species of glycosylated *Sc*Sec71. **b**, A schematic of the cryo-EM analysis of the chimeric Sec complex in an apo state. **c** and **d**, Distributions of particle view orientations in the final reconstructions of Classes 1 (c) and 2 (d). **e** and **f**, Fourier shell correlation (FSC) curves and local resolution maps of the final reconstructions. **g**, Superimposition of the Class 1 and 2 atomic models (based on the cytosolic domains) shows a slight difference in relative positions between Sec63-Sec71-Sec72 and the Sec61 complex. **h**, Side views showing the contact between the engineered cytosolic loops of Sec61α and the FN3 domain of *Sc*Sec63. Note that in Apo Class 2, the contact is more poorly packed than Class 1.

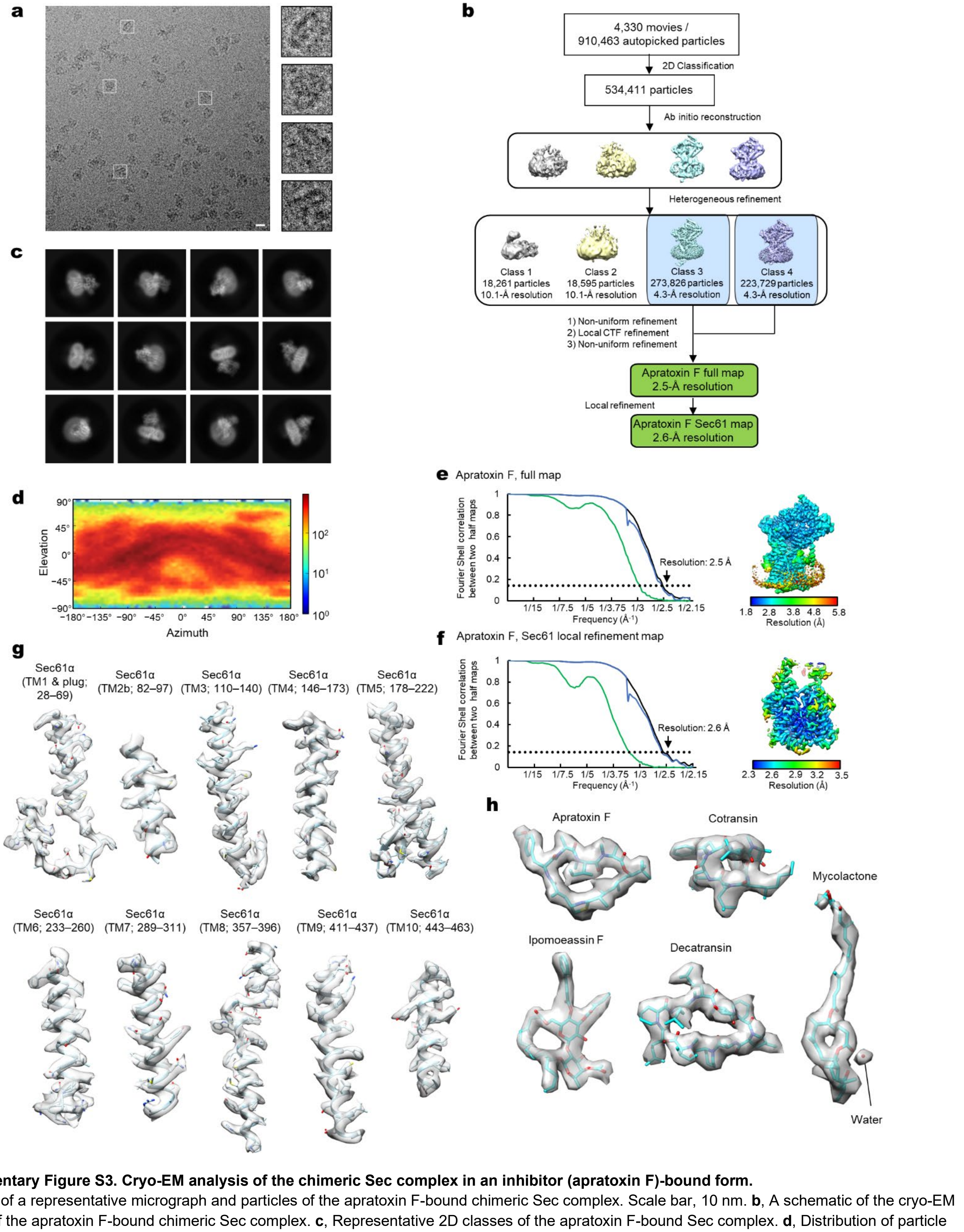


**Supplementary Figure S3. Cryo-EM analysis of the chimeric Sec complex in an inhibitor (apratoxin F)-bound form.**
**a**, Images of a representative micrograph and particles of the apratoxin F-bound chimeric Sec complex. Scale bar, 10 nm. **b**, A schematic of the cryo-EM analysis of the apratoxin F-bound chimeric Sec complex. **c**, Representative 2D classes of the apratoxin F-bound Sec complex. **d**, Distribution of particle view orientations in the final reconstruction. **e**, The FSC curve and local resolution map of the final reconstruction (full Sec complex map). **f**, As in e, but for the map from focused (local) refinement. **g**, Segmented density maps of the apratoxin F-bound Sec61α subunit. **h**, Segmented density features of bound natural inhibitors.

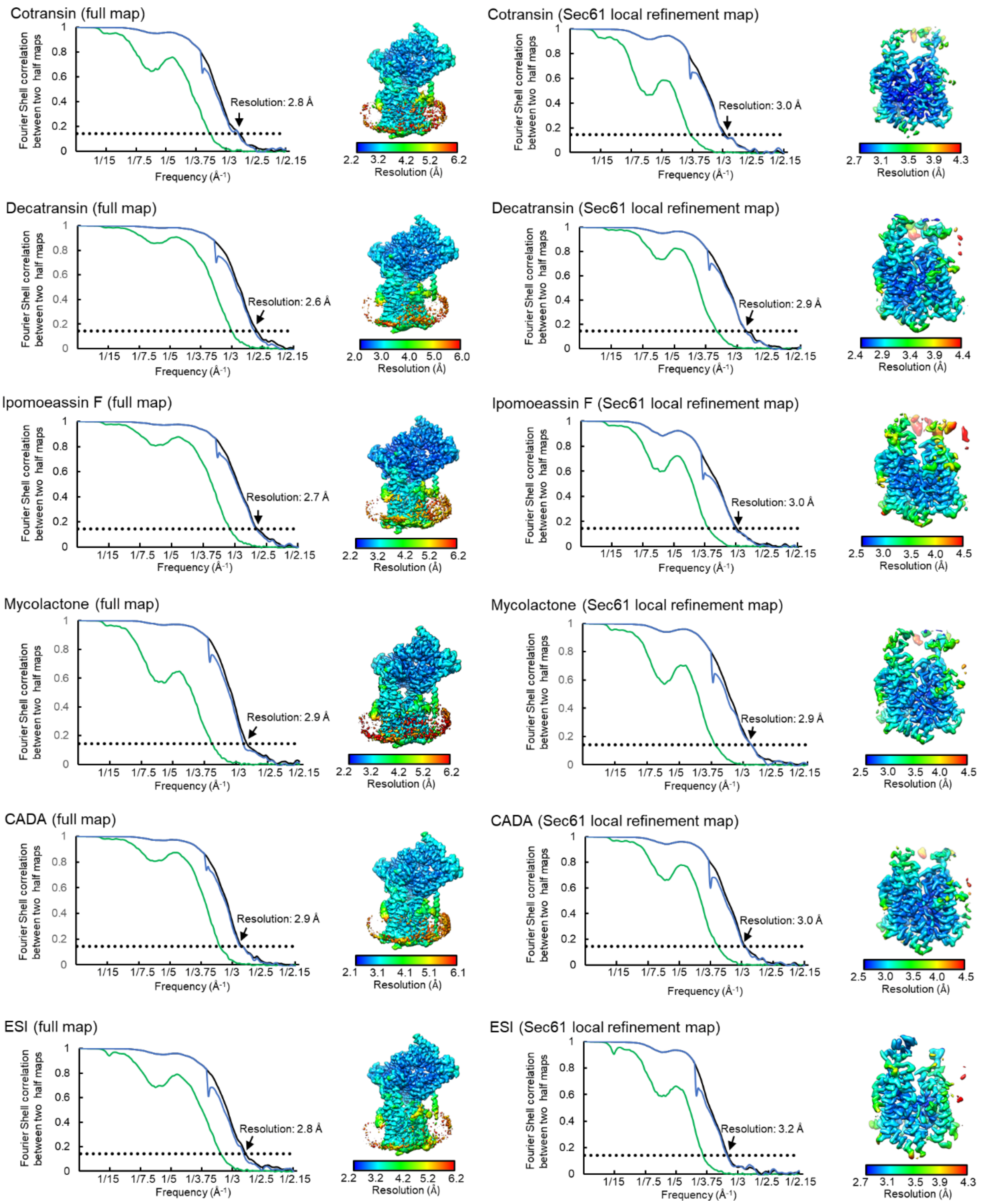


**Supplementary Figure S4. FSC curve and local resolution maps of inhibitor-bound Sec complexes.**
As in Supplementary Figure S3 e and f, but for all other inhibitor-bound structures.

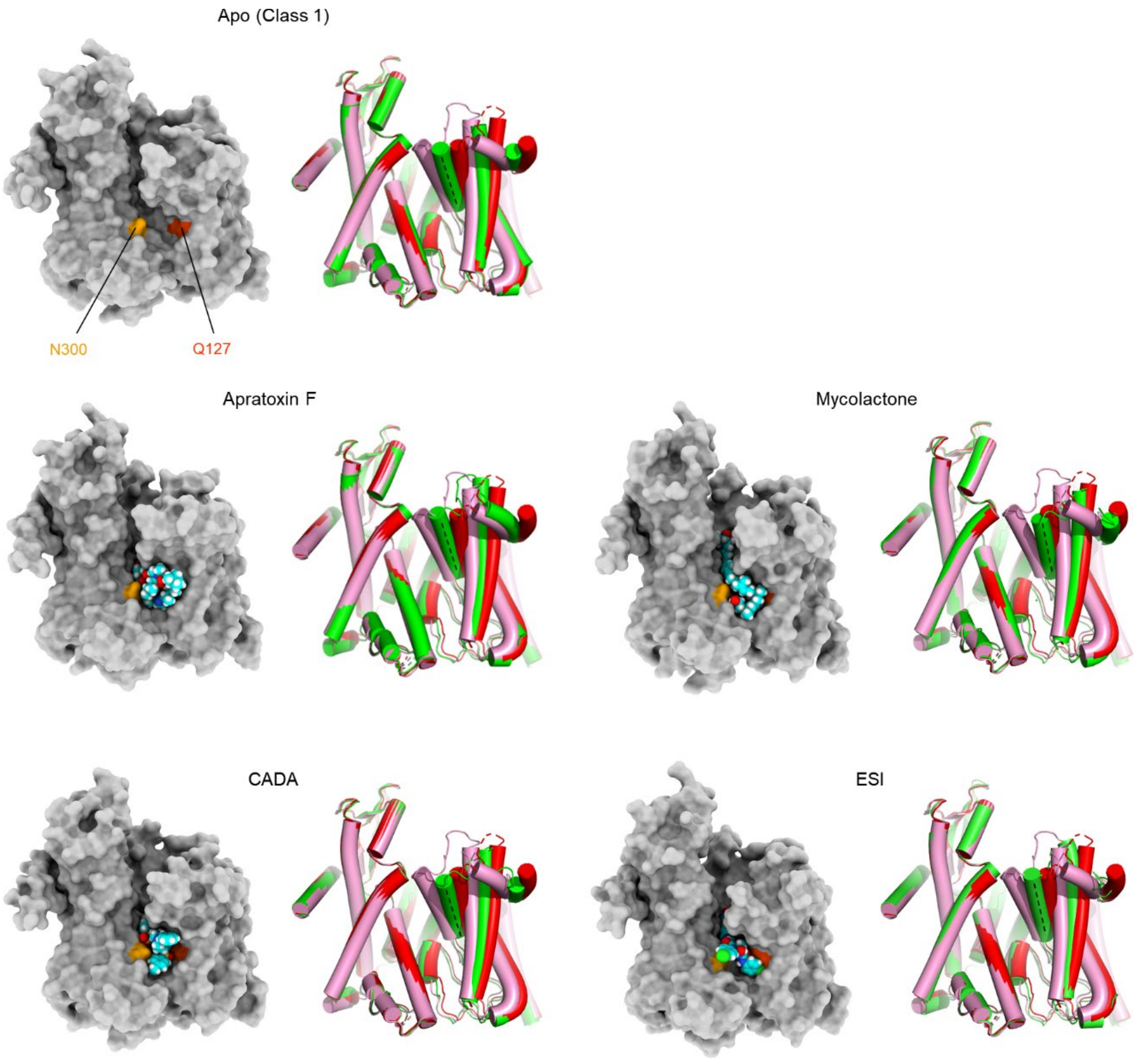


**Supplementary Figure S5. Variation in the extent of lateral gate opening in inhibitor-bound structures.**
As in Fig. 2 a and b, but showing other inhibitor-bound structures. In all panels showing a lateral gate comparison, cylindrical representations in red and pink are the cotransin- and ipomoeassin F- bound structures, respectively, whereas the representation in green is the structure with the indicated inhibitor.

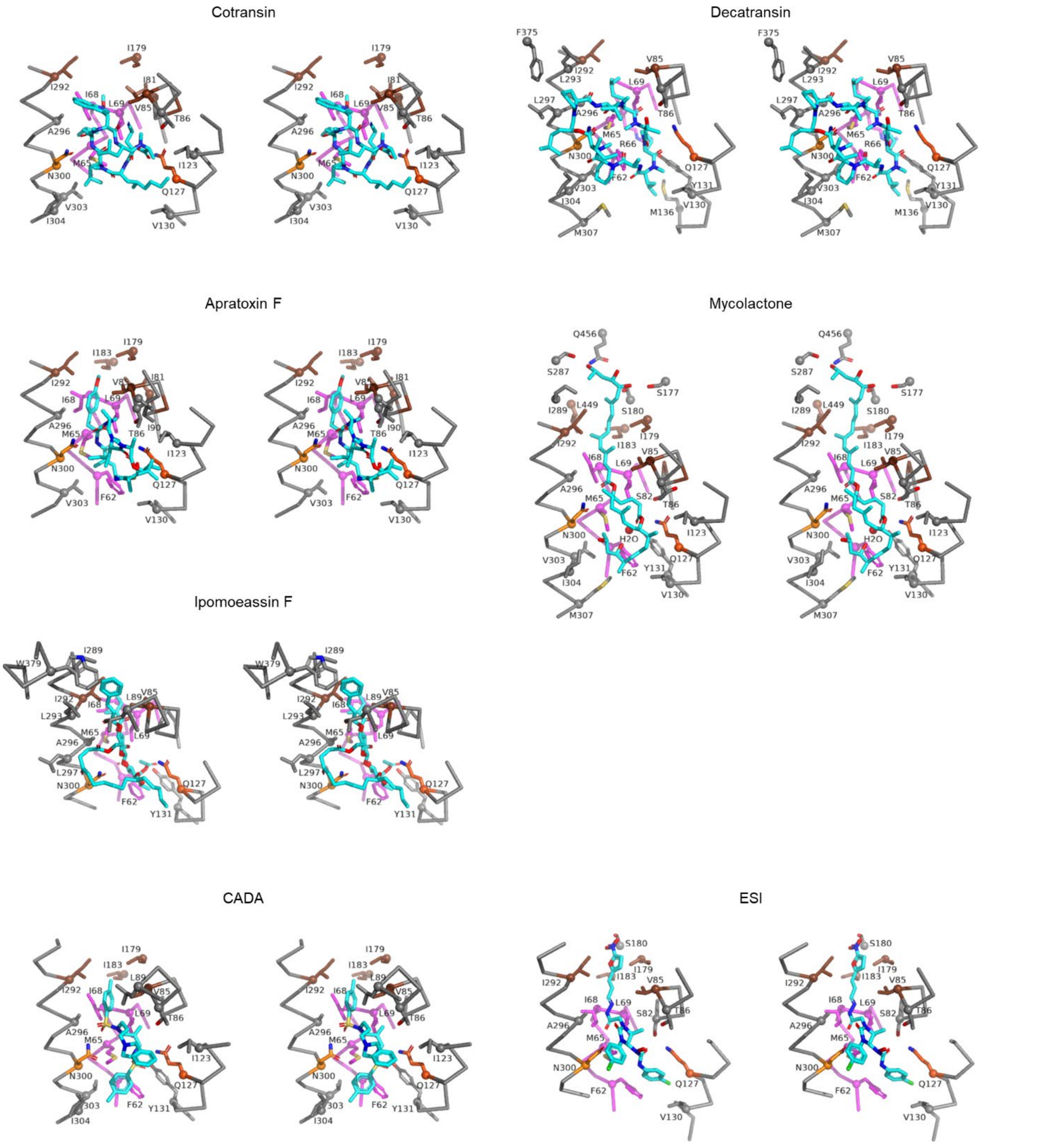


**Supplementary Figure S6. 3-D maps for interactions between Sec61 and inhibitors.**
Shown are stereo-views into the inhibitor-binding site. Inhibitors and adjacent protein side chains are shown in a stick representation together with Cα traces for TM2b, TM3, TM7, and the plug. The views are roughly similar between the different structures but adjusted for each structure for more clear representations. The following colors are used to differentiate parts: brown, pore ring residues; magenta, plug; lighter orange; N300, darker orange, Q127. All inhibitors are shown in cyan with certain atom-dependent coloring (nitrogen-blue, oxygen-red, sulfur-yellow, and chlorine-green).

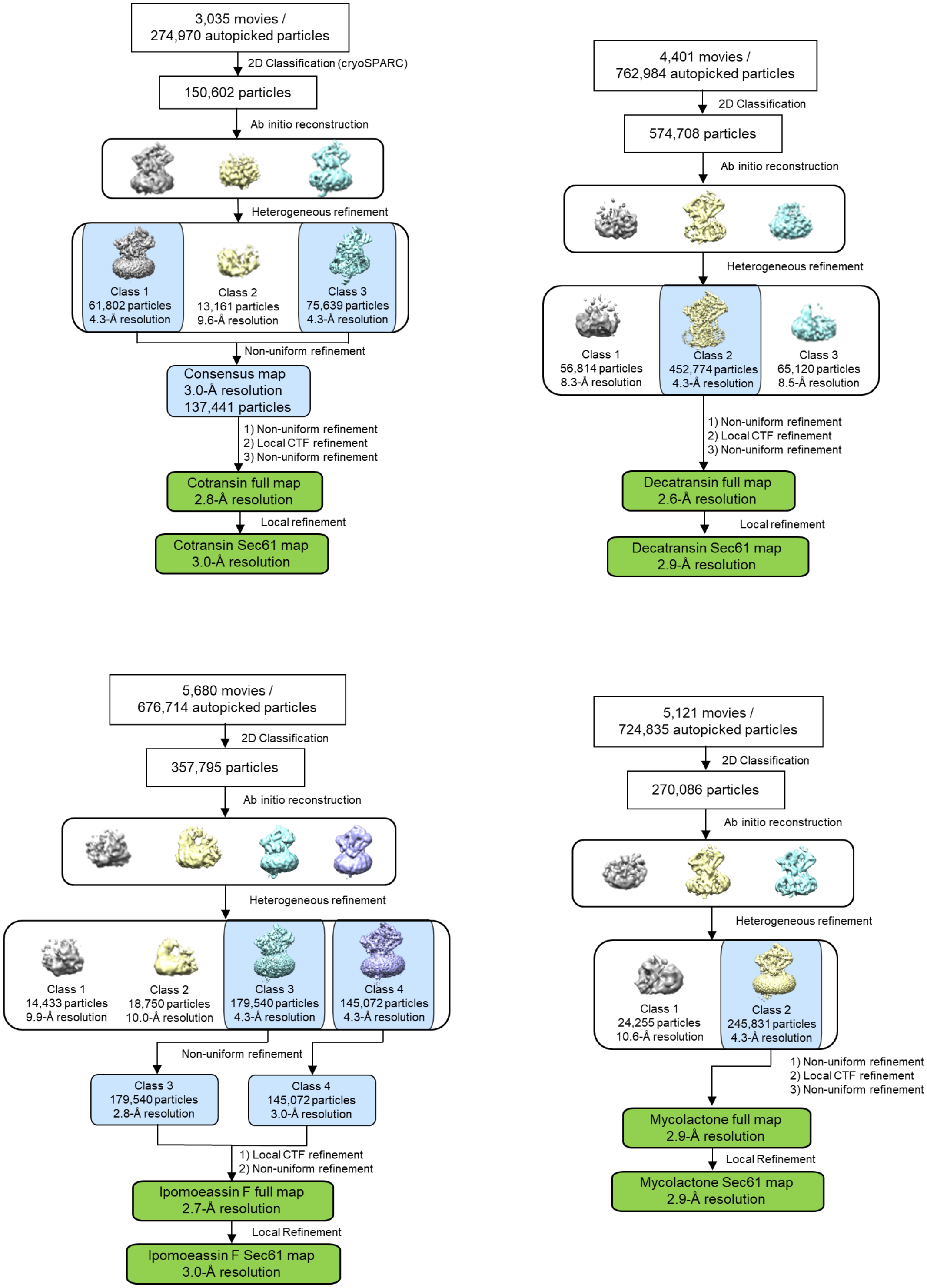


**Supplementary Fig. S7. Additional schematics of single-particle cryo-EM image analysis workflow.**
As in Supplementary Fig. S4b, but showing the datasets for the chimeric Sec complex in association with cotransin, decatransin, ipomoeassin F, or mycolactone.

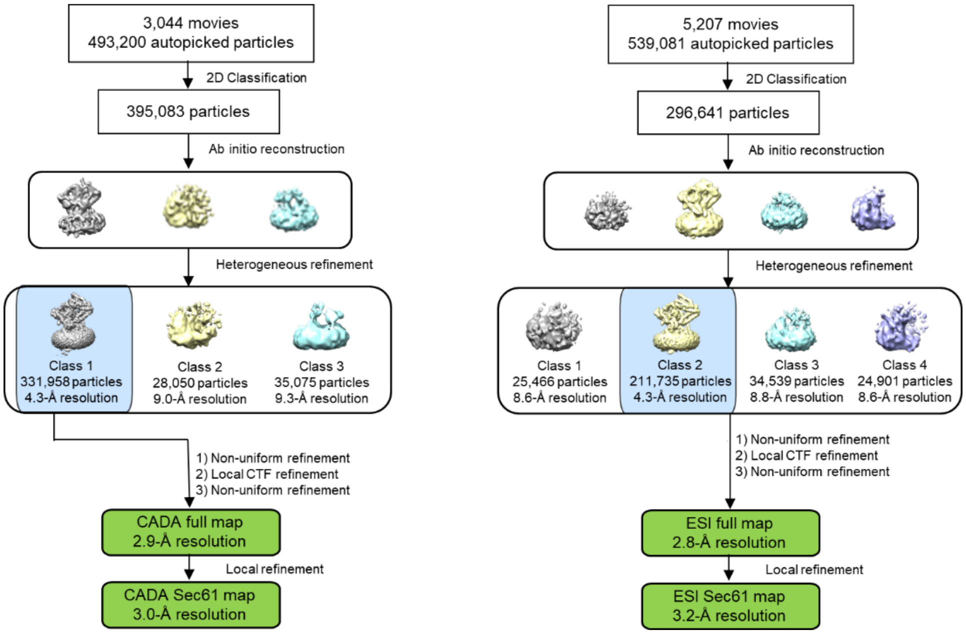


**Supplementary Fig. S8. Additional schematics of single-particle cryo-EM image analysis workflow.**
As in Supplementary Fig. S4b, but showing the datasets for the chimeric Sec complexes in association with CADA or ESI.

**Supplementary Table S1. Cryo-EM data collection, refinement and validation statistics**

| | Sec61, Apo (class 1) | Sec61, Apo (class 2) | Sec61, Cotransin | Sec61, Decatransin | Sec61, Apratoxin F | Sec61, Mycolactone | Sec61, Ipomoeassin F | Sec61, CADA | Sec61, ESI |
|---|---|---|---|---|---|---|---|---|---|
| PDB accession ID | 8DNV | 8DNW | 8DNX | 8DNY | 8DNZ | 8DO0 | 8DO1 | 8DO2 | 8DO3 |
| EMDB accession number | 27581 | 27582 | 27583 | 27584 | 27585 | 27586 | 27587 | 27588 | 27589 |
| **Data collection and processing** | | | | | | | | | |
| Magnification | 81,000x | 81,000x | 81,000x | 81,000x | 81,000x | 81,000x | 81,000x | 81,000x | 81,000x |
| Voltage (kV) | 300 | 300 | 300 | 300 | 300 | 300 | 300 | 300 | 300 |
| Electron exposure ($e^{-}$/Å$^2$) | 50 | 50 | 50 | 50 | 50 | 50 | 50 | 50 | 50 |
| Defocus range (μm) | -0.8 to -1.6 | -0.8 to -1.6 | -0.8 to -1.6 | -0.8 to -1.6 | -0.8 to -1.6 | -0.8 to -1.6 | -0.8 to -1.6 | -0.8 to -1.6 | -0.8 to -1.6 |
| Pixel size (Å) | 1.05 | 1.05 | 1.05 | 1.05 | 1.05 | 1.05 | 1.05 | 1.05 | 1.05 |
| Symmetry imposed | *C*1 | *C*1 | *C*1 | *C*1 | *C*1 | *C*1 | *C*1 | *C*1 | *C*1 |
| Initial particle images (no.) | 616,121 | 616,121 | 274,970 | 762,984 | 910,463 | 724,835 | 676,714 | 493,200 | 539,081 |
| Final particle images (no.) | 188,637 | 147,081 | 137,441 | 452,774 | 497,555 | 245,831 | 324,612 | 331,958 | 211,735 |
| Map resolution (Å) | 3.0 | 3.4 | 3.0 | 2.9 | 2.6 | 2.9 | 3.0 | 3.0 | 3.2 |
| FSC threshold | 0.143 | 0.143 | 0.143 | 0.143 | 0.143 | 0.143 | 0.143 | 0.143 | 0.143 |
| Map resolution range (Å) (min. to 75 percentile) | 2.6–7.4 | 2.9–7.5 | 2.6–7.6 | 2.4–6.4 | 2.3–6.4 | 2.6–7.5 | 2.6–7.6 | 2.3–7.2 | 2.7–6.9 |
| **Refinement** | | | | | | | | | |
| Initial model used | PDB:8DNZ | PDB:8DNZ | PDB:7KAH | PDB:8DNZ | PDB:8DNX | PDB:8DNZ | PDB:8DNZ | PDB:8DNZ | PDB:8DNZ |
| Model resolution (Å) | 3.2 | 3.5 | 3.1 | 3.0 | 2.7 | 3.1 | 3.2 | 3.1 | 3.4 |
| FSC threshold | 0.5 | 0.5 | 0.5 | 0.5 | 0.5 | 0.5 | 0.5 | 0.5 | 0.5 |
| Map sharpening *B* factor (Å$^2$) | -91 | -100 | -78 | -100 | -77 | -90 | -99 | -96 | -104 |
| Model composition | | | | | | | | | |
| Non-hydrogen atoms | 4157 | 4060 | 4,271 | 4,331 | 4,337 | 4,247 | 4,339 | 4,226 | 4,185 |
| Protein residues | 535 | 528 | 543 | 549 | 551 | 542 | 551 | 541 | 534 |
| Ligands | 0 | 0 | 1 | 1 | 1 | 1 | 1 | 1 | 1 |
| *B* factors (Å$^2$) | | | | | | | | | |
| Protein | 76.57 | 49.58 | 71.46 | 41.25 | 63.60 | 52.92 | 65.58 | 49.43 | 61.03 |
| Ligand | - | - | 60.88 | 39.34 | 56.47 | 42.82 | 47.33 | 47.21 | 62.00 |
| R.m.s. deviations | | | | | | | | | |
| Bond lengths (Å) | 0.004 | 0.003 | 0.004 | 0.003 | 0.004 | 0.002 | 0.003 | 0.004 | 0.004 |
| Bond angles (°) | 0.593 | 0.515 | 0.579 | 0.597 | 0.513 | 0.489 | 0.560 | 0.650 | 0.582 |
| **Validation** | | | | | | | | | |
| MolProbity score | 1.51 | 1.30 | 1.47 | 1.47 | 1.32 | 1.40 | 1.67 | 1.38 | 1.43 |
| Clashscore | 7.53 | 5.57 | 6.35 | 7.65 | 5.90 | 7.31 | 10.10 | 6.98 | 6.70 |
| Poor rotamers (%) | 0 | 0 | 0 | 0 | 0 | 0.22 | 0.22 | 0.22 | 0 |
| Ramachandran plot | | | | | | | | | |
| Favored (%) | 0 | 0 | 0 | 0 | 0 | 0 | 0 | 0 | 0 |
| Allowed (%) | 2.48 | 1.36 | 2.63 | 2.22 | 1.29 | 1.69 | 2.76 | 1.69 | 2.30 |
| Disallowed (%) | 97.52 | 98.64 | 97.37 | 97.78 | 98.71 | 98.31 | 97.24 | 98.31 | 97.70 |

**Supplementary Table S2. Effects of mutations in *Sc*Sec61 on yeast growth inhibition by cotransin, ipomoeassin F, and decatransin**

| | | | IC50 value (μM) | | |
|---|---|---|---|---|---|
| ScSec61 aa position | Mutation | Position in HsSec61A1 | Cotransin | Ipomoeassin F | Deca-transin* |
| | WT | | 0.87; 0.55[#] | 0.06 | 3.1; 1.2* |
| 47 | G47D | C46 | >200* | | 100* |
| 63 | L63D | F62 | >200 | 0.40 | |
| 63 | L63W | F62 | 0.43 | 0.25 | |
| 63 | L63N | F62 | >200 | <0.06 | 2.9* |
| 71 | A71D | A70 | 1.4* | | 3* |
| 72 | S72F | S71 | >200 | 0.17 | >200* |
| 79 | E79K | E78 | >200* | | >200* |
| 81 | G81D | G80 | >200* | | 3.6* |
| 82 | V82D | I81 | >200 | >100 | |
| 82 | V82W | I81 | >200 | | |
| 84 | P84L | P83 | >200* | | >200* |
| 86 | I86T | V85 | 0.54 | <0.06 | |
| 86 | I86D | V85 | >200 | <0.06 | |
| 86 | I86W | V85 | 2.08 | | |
| 87 | T87I | T86 | >200* | | >200* |
| 89 | S89D | G88 | 1.10 | <0.06 | |
| 89 | S89W | G88 | 1.00 | 0.09 | |
| 90 | M90D | L89 | >200 | >100 | |
| 90 | M90W | L89 | 1.06 | 0.13 | |
| 93 | Q93D | Q92 | 2.72 | 0.07 | |
| 93 | Q93W | Q92 | 0.53 | 0.12 | |
| 96 | Q96D | A95 | 0.77 | <0.06 | |
| 96 | Q96W | A95 | 1.10 | 0.07 | |
| 97 | G97D | G96 | 0.62; 0.3* | 1.48 | >200* |
| 97 | G97W | G96 | 1.04 | 0.06 | |
| 111 | R111D | R109 | 1.15 | <0.06 | |
| 111 | R111W | R109 | 0.87 | <0.06 | |
| 115 | Q115D | N113 | 1.84 | <0.06 | |
| 115 | Q115W | N113 | 1.05 | <0.06 | |
| 129 | Q129D | Q127 | >200 | 0.28 | |
| 129 | Q129W | Q127 | >200 | >100 | |
| 129 | Q129L | Q127 | 0.58; 1.1[#] | 18.3 | >100 |
| 168 | D168W | D166 | 0.22 | <0.06 | |
| 172 | S172D | Q170 | 1.09 | <0.06 | |
| 172 | S172W | Q170 | 0.88 | 0.07 | |
| 178 | G178D | G176 | 1.18 | <0.06 | |
| 178 | G178W | G176 | 0.52 | <0.06 | |
| 179 | S179D | S177 | 1.03 | <0.06 | |
| 179 | S179W | S177 | 0.26 | <0.06 | |
| 179 | S179A | S177 | 0.82 | | |
| 179 | S179C | S177 | 0.68 | | |
| 179 | S179F | S177 | 0.56 | | |
| 179 | S179G | S177 | 0.80 | | |
| 179 | S179H | S177 | 1.00 | | |
| 179 | S179I | S177 | 0.53 | | |
| 179 | S179K | S177 | 0.57 | | |
| 179 | S179L | S177 | 1.03 | | |
| 179 | S179M | S177 | 1.01 | | |
| 179 | S179N | S177 | 1.04 | | |
| 179 | S179Q | S177 | 1.13 | | |
| 179 | S179R | S177 | 0.54 | | |
| 179 | S179S(=WT) | S177 | 0.91 | | |
| 179 | S179T | S177 | 0.98 | | |
| 179 | S179V | S177 | 0.80 | | |
| 179 | S179Y | S177 | 0.56 | | |
| 181 | I181D | I179 | 0.59 | <0.06 | |
| 181 | I181W | I179 | <0.1 | | |

| | | | IC50 value (μM) | | |
|---|---|---|---|---|---|
| ScSec61 aa position | Mutation | Position in HsSec61A1 | Cotransin | Ipomoeassin F | Deca-transin* |
| 182 | S182D | S180 | 1.34 | <0.06 | |
| 182 | S182W | S180 | >200 | 0.06 | |
| 185 | T185D | I183 | >200 | >100 | |
| 185 | T185W | I183 | 0.60 | | |
| 186 | A186T | A184 | 0.3* | | 2.4* |
| 287 | Y287D | Y285 | >200 | | |
| 287 | Y287W | Y285 | 5.32 | | |
| 291 | T291W | I289 | 0.83 | 13.0 | |
| 291 | T291D | I289 | 3.05 | 0.07 | |
| 294 | M294D | I292 | >200 | >100 | |
| 294 | M294W | I292 | 1.06 | | |
| 296 | Q296D | Q294 | 1.07 | 0.14 | |
| 296 | Q296W | Q294 | 0.59 | <0.06 | |
| 298 | A298T | A296 | >200* | | >200* |
| 302 | N302L | N300 | >200 | >100 | >100 |
| 302 | N302D | N300 | 0.51 | >100 | |
| 302 | N302W | N300 | >200 | >100 | |
| 305 | L305D | V303 | >200 | 11.7 | |
| 305 | L305W | V303 | >200 | <0.06 | |
| 307 | S307D | S305 | 1.37 | <0.06 | |
| 307 | S307W | S305 | >200 | 0.58 | |
| 307 | S307F | S305 | >200* | | >200* |
| 379 | T379D | T378 | 1.07 | 0.09 | |
| 379 | T379W | T378 | 0.59 | <0.06 | |
| 380 | W380D | W379 | 1.07 | 0.15 | |
| 382 | E382D | E381 | 1.09 | <0.06 | |
| 382 | E382W | E381 | 1.70 | <0.06 | |
| 384 | S384D | S383 | 2.01 | <0.06 | |
| 384 | S384W | S383 | 0.60 | 0.19 | |
| 386 | T386D | S385 | 1.44 | <0.06 | |
| 386 | T386W | S385 | 1.11 | <0.06 | |
| 430 | G430D | G429 | 0.5* | | 3.7* |
| 446 | A446T | T445 | 1.3* | | 2.7* |
| 450 | M450D | L449 | 4.04 | 0.87 | |
| 450 | M450W | L449 | 2.15 | | |
| 454 | T454D | I453 | 0.46 | <0.06 | |
| 454 | T454W | I453 | <0.1 | <0.06 | |
| 461 | A461D | I460 | 1.36 | | |
| 461 | A461W | I460 | 0.86 | | |
| 480 | M480D | L475 | 1.01 | | |
| 480 | M480W | L475 | 1.03 | | |
| multiple | ΔPlug (52-74→G) | | >200 | >100 | |
| multiple | V82D/I86D/M294K | I81/V85/I292 | >200 | | |
| multiple | V82D/I86D/M450K | I81/V85/L449 | >200 | | |
| multiple | I181D/T185D/M450K | I179/I183/L499 | >200 | | |
| multiple | Q308/I323/W326/L342A | Q306/L321/W324/L341 | >200 | | |
| multiple | I86T/Q308/I323/W326/L342A | V85/Q306/L321/W324/L341 | >200 | | |
| multiple | Q96W/Q99H | A95/K98 | 0.89 | | |

Gray highlight: IC50 larger than 5x but less than 100x of IC50 of WT.
Yellow highlight: IC50 larger than 100x of IC50 of WT.
Blank: not determined.
* Data from Junne et al., doi:10.1242/jcs.165746.
[#] Values measured with the strain BY4743Δ9aURA harboring pDQ1.
